# A Conformational Switch in the Catalytic Lysine Governs the Stereochemical Outcome of DERA-Catalyzed Aldol Addition

**DOI:** 10.64898/2026.07.26.740800

**Authors:** Srita Dutta, Ananya Nayak, Kodru Jeevani, Thangavelu Saravanan, Jagannath Mondal, Anand T. Vaidya

## Abstract

2-Deoxyribose-5-Phosphate Aldolase (DERA) is a key enzyme in the pentose phosphate pathway. Due to its C-C bond formation and stereoselective capabilities, DERA has been widely used for biocatalytic applications including the synthesis of chiral intermediates for antiviral and anticancer drugs. Protein engineering has expanded its substrate-scope, improved yield, and enhanced *S*-stereoselectivity. Recently, rational mutagenesis of DERA from *Geobacillus sp.* (DERA*_Gs_*) resulted in two enantiocomplementary variants, T12I/S185A and S185G. In particular, the single mutation at Ser185 to Gly inverts the natural *S*-selectivity to *R*-selectivity. However, the molecular basis of enantioselectivity remains unclear. Here, we determined high-resolution crystal structures of wild-type DERA*_Gs_* and two enantiocomplementary variants in the apo form. Using a combination of structural biology, biochemistry, biocatalysis and molecular dynamic simulations, we show that the catalytic lysine adopts two conformations, a finding that relates to the enantioselectivity of DERA. We further identified a mechanism of regulating enantioselectivity via a key conserved amino acid. Using DERA from *E. coli*, we show that these findings are conserved across bacterial species.

## Introduction

The formation of new carbon-carbon bonds with precise stereochemistry is a central component of synthetic chemistry and pharmaceuticals.^1–4^ Nature achieves this via several enzymes that synthesize key intermediates in biosynthetic pathways in the cell. The 2-deoxyribose-5-phosphate aldolase (DERA) is one such enzyme that catalyses the reversible aldol addition of acetaldehyde and D-glyceraldehyde-3-phosphate, resulting in 2-deoxy-D-ribose-5-phosphate.^5–8^ DERAs have been engineered (i) to accommodate a variety of donor substrates, such as acetone, hydroxyacetone, dihydroxyacetone, fluoroacetone, butanone, cyclobutanone, cyclopentanone, propanal, glycolaldehyde, etc., and a variety of aliphatic, aryl and heteroaryl aldehydes as acceptor substrates,^4,9,10,13,15^ (ii) to increase the yields of the non-natural substrates,^9,11–13^ and (iii) to reverse the enantioselectivity.^9,20^ Given this versatility, DERAs are widely used for biocatalytic applications such as, in the synthesis of anticancer agents like epothilones, nucleotide analogs for antivirals like Islatravir, a diverse range of deoxy sugars, pyrimidine nucleosides, etc.^14,16,17^ DERAs have also been used to catalyse sequential aldol addition reactions, to generate the precursor for statin (atorvastatin, rosuvastatin, and compactin) synthesis, and in combination with other enzymes to generate various chiral centres.^18,19^ While the mechanism of aldol addition of Class I aldolases such as DERA, involving an enamine intermediate, is well characterized, the molecular/atomic details underlying the enantioselectivity and its regulation are still unclear.

Bisterfeld et al. used molecular dynamics (MD) simulations and homologous grafting on the enantiocomplementary aldolase enzymes 2-keto-3-deoxy-6-phosphogluconate aldolase (KDPG) and 2-keto-3-deoxy-6-phosphogalactonate aldolase (KDPGal), to identify the key residues responsible for their stereoinversion. ^20^ Structural alignment using KDPG, KDPGal and DERA from *E.coli* (DERA*_Ec_*) followed by aldol screening using acetaldehyde and propanal, identified T18 and A203 of DERA*_Ec_* as key residues. Mutations at these positions resulted in decreased enantiomeric excess (T18S: 20% ee, A203G: 88% ee) compared to wild-type DERA*_Ec_* (∼96% ee).^20^ Recently, Saravanan and coworkers rationally engineered DERA from *Geobacillus sp.* (DERA*_Gs_*) to accept a variety of heteroaryl aldehydes as acceptors and acetone as the donor (Fig 1A). Targeting positions T12 and S185 (corresponding to T18 and A203 of DERA*_Ec_*), they identified two enantiocomplementary variants: the double mutant, T12I/S185A, which exhibited enhanced *S*-selectivity (*S*/*R* = 97:3 e.r.) compared with the wild-type (*S*/*R* = 77:23 e.r.), and the single mutant, S185G, which showed inverted *R*-selectivity (*S*/*R* = 11:89 e.r.).^9^ Despite the key roles of these residues of DERA in the enantioselectivity of aliphatic and aryl aldol products, the molecular details and rationale are lacking.

**Figure 1.**
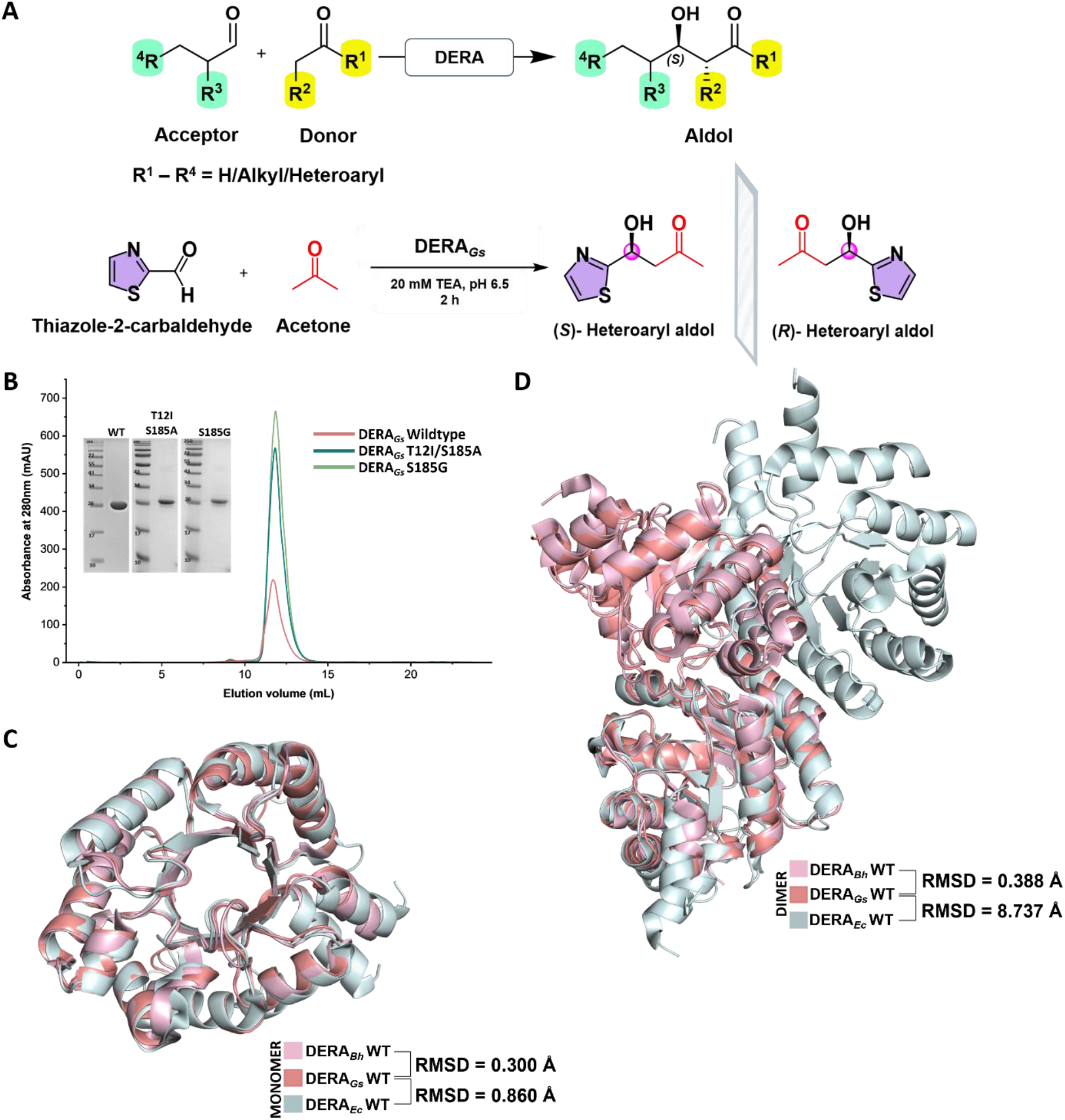
A structural comparison of DERA_Gs_ with its homologs: (A) A general chemical scheme of DERA-catalyzed aldol addition reactions using non-natural acceptor and donor substrates (above) and the chemical scheme for the cross-aldol addition of thiazole-2-carbaldehyde and acetone catalysed by DERA*_Gs_* (below). (B) Size exclusion chromatogram overlay of different DERA*_Gs_* samples and their SDS-PAGE analysis. (C) The protein adopts the characteristic triosephosphate isomerase (TIM) barrel fold conserved among DERA enzymes. Superposition of monomeric DERA*_Gs_* with DERA*_Bh_* and DERA*_Ec_* shows a conserved overall architecture. RMSD values show the conserved structure of the three enzymes. (D) DERA*_Gs_* and DERA*_Bh_* have a similar dimer, which is different form the DERA*_Ec_,* as shown by the RMSD and the structural superposition.

Here, we determine three crystal structures of DERA*_Gs_*-the wild-type, the T12I/S185A variant and the S185G variant, which provide atomic-level details of enantioselectivity in DERA. Molecular dynamic simulations of the apo and acetone (donor substrate) bound complexes corroborate the hypothesis and provide a possible mechanism. Finally, we show that analogous mutations in DERA*_Ec_*also generate the expected enantioselective products, indicating a conserved mechanism of enantioselectivity across different bacterial species.^21^

## Results

### Crystal structure of DERA*_Gs_* and its comparison to other DERAs

The crystal structures of DERA*_Gs_* wild-type (WT), the T12I/S185A variant and the S185G variant were determined at 1.19 Å, 1.22 Å and 1.50 Å resolutions, respectively (Table S1). The structure of DERA*_Bh_* (DERA from *Bacillus halodurans*, PDB ID: 6D33) was used as a model for molecular replacement as it has a 74% sequence identity with DERA*_Gs_*. The DERA*_Gs_* proteins were purified as dimers and the asymmetric unit has a dimer (Fig 1B) (Fig S1). The structures showed the typical TIM barrel fold observed in other DERA enzymes and the last 5-7 residues at the C-terminus could not be modelled due to weak electron density, likely because of flexibility (Fig 1C).^22^ The monomeric structure of DERA*_Gs_* is similar to DERA*_Bh_* and DERA*_Ec_*, with RMSD values of 0.3 Å and 0.9 Å, respectively, compared to DERA*_Gs_* WT, calculated over the Cα atoms (Fig 1C). Citrate ion(s), from the crystallization condition, is modelled in the wild-type and the S185G variant (Fig S2).

The DERA*_Gs_* dimer is similar to that of DERA*_Bh_*dimer with a RMSD of 0.4 Å, but it differs considerably from the DERA*_Ec_*dimer, with a RMSD of 8.7 Å (Fig 1E). A closer look at the dimer interface shows a larger dimer interface for DERA*_Gs_*involving the helices ɑ2, ɑ3, ɑ4, ɑ5, ɑ6 & ɑ7, and the loops 2, 4, 6, 8, 9 & 14 compared to the DERA*_Ec_* dimer that involves helices ɑ2, ɑ3 & ɑ4 and the loops 3 and 8 (Fig S3). This is reflected in the larger solvent buried surface area of 1277 Å^2^ in DERA*_Gs_* compared to 595 Å^2^ of DERA*_Ec_*. This could contribute to the higher thermostability of DERA*_Gs_* (T_m_ of 81.0 °C) compared to DERA*_Ec_* (T_m_ of 74.4 °C) (Fig S4). Comparing the DERA structures from eleven different organisms shows that seven of these are similar to the DERA*_Gs_* dimer, with the RMSD ranging from 0.39 Å to 0.98 Å. The remaining three have different dimers interfaces and show an arrangement similar to the DERA*_Ec_* dimer (Table 1).^6,7,23–25^ Taken together, the DERA*_Gs_*-like dimer has a larger solvent buried surface area compared to DERA*_Ec_* and is adopted by majority of the thermophiles.

**Table 1.** Classification of DERA structures based on the dimer interface. : The DERA structures from eleven organisms are grouped as either DERA*_Gs_*-like or DERA*_Ec_*-like based on the structural alignment with DERA*_Gs_* and DERA*_Ec_* dimers. In case of the structures involving tetramer, hexamer and 12-mer, all the possible dimer interfaces were aligned with the DERA*_Gs_* dimer and DERA*_Ec_* dimer, and classified accordingly.

| **S.No.** | **PDB ID** | **Organism** | **RMSD (Å)<br>Compared to DERA<sub>*Gs*</sub>** | **RMSD (Å)<br>Compared to DERA<sub>*Ec*</sub>** | **Category** | **Oligomeric State** | **Classification** |
|---|---|---|---|---|---|---|---|
| 1 | **44HZ**<br>(*current study*) | *Geobacillus sp.* | -- | 8.825 | Thermophile | Dimer | -- |
| 2 | **5DBT** | *Streptococcus suis* | 0.420 | 9.031 | Mesophile | 12-mer | **DERA<sub>*Gs*</sub>- like** |
| 3 | **6D33** | *Bacillus halodurans* | 0.343 | 8.674 | Mesophile | Hexamer | **DERA<sub>*Gs*</sub>- like** |
| 4 | **3R13** | *Thermotoga maritima* | 0.443 | 9.080 | Thermophile | Dimer | **DERA<sub>*Gs*</sub>- like** |
| 5 | **1J2W** | *Thermus thermophilus* | 0.553 | 9.447 | Thermophile | Tetramer | **DERA<sub>*Gs*</sub>- like** |
| 6 | **4XBS** | *Levilactobacillus brevis* | 0.760 | 8.882 | Mesophile | Dimer | **DERA<sub>*Gs*</sub>- like** |
| 7 | **3NG3** | *Mycobacterium avium* | 0.948 | 8.786 | Mesophile | Tetramer | **DERA<sub>*Gs*</sub>- like** |
| 8 | **1VCV** | *Pyrobaculum aerophilum* | 0.867 | 9.281 | Thermophile | Dimer | **DERA<sub>*Gs*</sub>- like** |
| 9 | **1P1X** | *Escherichia coli* | 8.825 | -- | Mesophile | Dimer | -- |
| 10 | **5C6M** | *Shewanella helifaxensis* | 9.287 | 0.486 | Mesophile | Tetramer | **DERA<sub>*Ec*</sub>- like** |
| 11 | **5C2X** | *Colwellia psychrerythraea* | 9.508 | 1.105 | Psychrophile | Dimer | **DERA<sub>*Ec*</sub>- like** |

### Two conformations of the active site lysine residue correlate with enantioselectivity of DERA

Comparison of the active site by structural superposition of DERA*_Gs_* WT and DERA*_Ec_* show that nearly all the residues involved in substrate binding and in catalysis adopt a similar conformation (Fig S5). However, in DERA*_Gs_* WT, the catalytic lysine residue (Lys154) adopts two conformations that differ by a 1.5 Å shift and a 63.5° rotation of the terminal amine nitrogen (Fig 2A, B). We name these conformations as conf-a and conf-b; the occupancy of conf-a is 56% and that for conf-b is 44% (Fig 2A). In both these conformations, the terminal amine nitrogen of Lys154 is involved in several hydrogen bonding interactions within the active site, but two hydrogen bonds, one with Asp91 and the other with Ser156, are mutually exclusive between the two conformations (Fig 2B, C). We use these two hydrogen bonds to define the two conformations - in conf-a the amine of Lys154 has a hydrogen bonding interaction with the carboxyl side chain of Asp91, while in conf-b the amine of Lys154 has a hydrogen bonding interaction with the main chain oxygen of Ser156 (Fig 2C). Previously reported structures of DERA, from different organisms, have the catalytic lysine in the conf-a, but the second conformation has not been reported in literature (Fig S6). The crystal structure of the two enantiocomplementary variants of DERA*_Gs_* reveal two conformations of the Lys154. In the DERA*_Gs_* T12I/S185A variant, which results in a *S*-isomer with 94% ee, the Lys154 is similar to conf-a, and in the DERA*_Gs_* S185G variant, which results in a *R*-isomer with 74% ee, the Lys154 is close to the conf-b (Fig 2A). The terminal amine nitrogen of Lys154 in the S185G variant is in a similar position in the conf-b of the WT, but the carbon side chain of Lys154 adopts a different conformation, and we term this as conf-c in the S185G variant. From a catalytic and enantioselectivity perspective, conf-b and conf-c are most likely equivalent as the terminal amine nitrogen is in a similar position and are involved in similar hydrogen bonding interactions, including with the Ser156 (Fig 3A, B, S7). Taken together, the catalytic Lys154 adopts two major conformations, with exclusive Hydrogen-bonds and these conformations corelate with the enantioselectivity: conf-a in the T12I/S185A variant (*S*/*R*=97:03 e.r.) and conf-c in S185G (*S*/*R*=11: 89 e.r.). DERA*_Gs_*wild-type adopts two conformations, conf-a and conf-b (similar to conf-c), which may account for its moderate enantioselectivity (*S*/*R*=77:23 e.r.).

**Figure 2.**
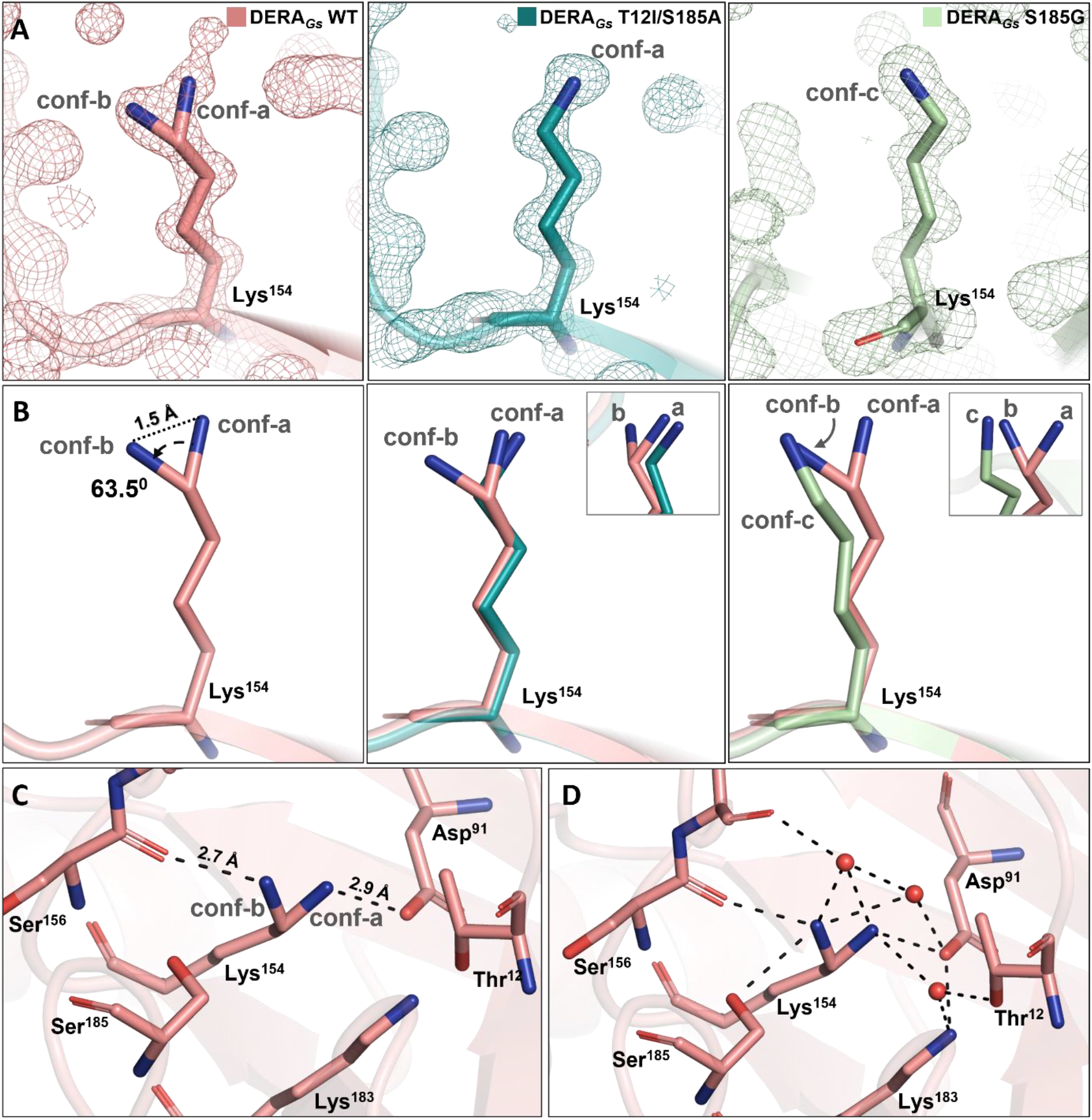
The catalytic lysine adopts different conformations, stabilized by H-bonding network: (A) The 2F_o_-F_c_ electron density maps contoured at 1.5σ, shows the alternate conformation of the catalytic Lys154 in the DERA*_Gs_* WT, and the single conformations in the T12I/S185A and S185G variants. In WT, the occupancy of conf-a & conf-b were 56% and 44% respectively, in monomer A and 51% and 49% in monomer B. (B) The terminal amine nitrogen of the Lys154 in the two conformations (conf-a, conf-b) of the DERA*_Gs_* WT are 1.5 Å and 63.5° away from each other. Superposition of the DERA*_Gs_* WT structure with the two variants reveal that the Lys154 in T12I/S185A adopts a conformation similar to conf-a, and the conf-c in S185G is similar to conf-b. The inset shows the superposition viewed from a different angle. (C) The side chain of Lys154 is stabilized by H-bonding to Ser156 in conf-b and to Asp91 in conf-a. (D) The solvent network stabilizing both the conformations of the catalytic lysine are shown. Active-site residues are shown as sticks, water molecules are depicted as red spheres, and H-bonds are represented by dashed lines.

**Figure 3.**
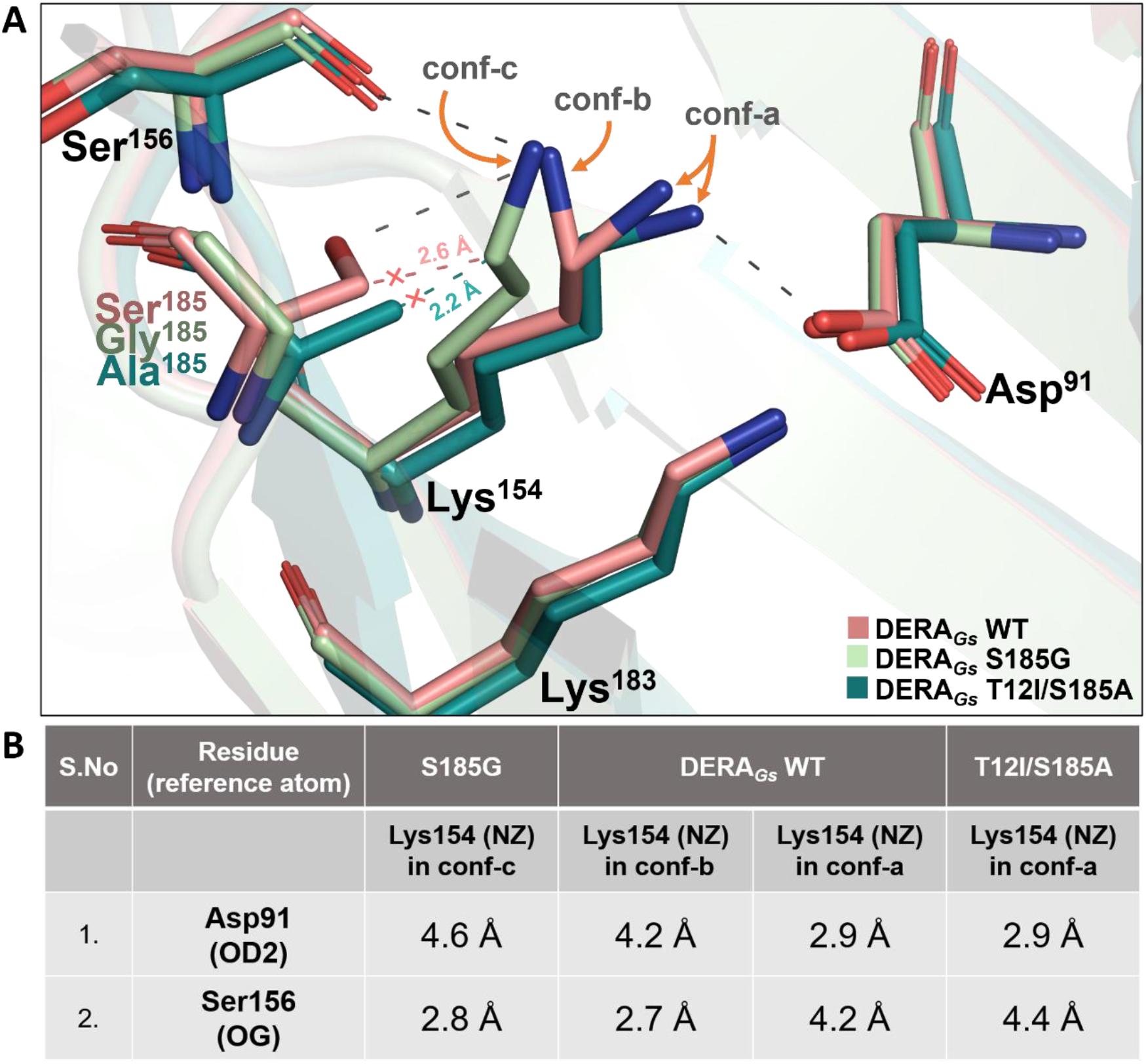
The role of residue 185 in regulating the conformation of Lys154 in DERA*_Gs_*: (A) The superposition of the DERA*_Gs_* WT and the two variants show that the S185G mutation creates a pocket, which accommodates the novel conf-c state of Lys154. In the T12I/S185A mutant, the alanine –CH_3_ group sterically hinders the ε-C atom of the Lys154 conf-c (shown in teal cross and distance), thus favoring conf-a. In the WT enzyme, Ser185 similarly disfavors conf-c (shown in salmon cross and distance), but the hydroxyl group of Ser185 forms a hydrogen bond with terminal amine nitrogen of Lys154. The hydrogen bonds shown as dashed lines (black) correspond to the DERA*_Gs_* wild-type structure. (B) The distances of the carboxylic side chain oxygen of Asp91(OD2) and the main chain oxygen of Ser156(OG) from the terminal nitrogen of Lys154 (NZ) in all three conformations are tabulated. Despite the different conformations of the Lys154 carbon chains in conf-b and conf-c, the hydrogen bonding with Ser156 is maintained which is unlike conf-a that hydrogen bonds with Asp91.

An overlay of the three structures reveals that the three conformations of the catalytic lysine are driven by a combination of steric effect and electrostatic interactions, involving the serine/alanine/glycine at 185 position. In the T12I/S185A variant, the Cβ carbon of the A185 would have a steric clash with the Cε carbon of the K154 in conf-c, so it adopts the conf-a (Fig 3A). In the S185G variant, due to the lack of a bulky sidechain in the G185 residue, it opens up a pocket that stabilizes the sidechain of K154 into conf-c. In case of the wild-type enzyme, opposing forces of steric clash and hydrogen bonding interactions between the S185 and the K154 results in stabilizing two conformations of the K154; a steric clash between Cβ carbon of the S185 and the Cε carbon of the K154 in the conf-c leads to conf-a, but the sidechain hydroxyl of S185 can form hydrogen bonding with the amine of the K154, resulting in an intermediate conformation conf-b. In conf-b and conf-c, the carbons of the lysine sidechain occupy different positions, but the terminal amine nitrogen that is involved in catalysis is in a similar position (Fig 3B). Thus, the conf-b and conf-c would be catalytically similar, but different from conf-a.

### Molecular dynamics simulations of the apo and donor bound DERA*_Gs_*reveal two conformations of Lys154

To corroborate the observations from the crystal structures and to ascertain the conformation of the Lys154 in the two enantiocomplementary variants, we performed molecular dynamics simulations of the two variants of DERA*_Gs_*. To avoid bias, the crystal structures of the S185G and T12I/S185A variants were not used for the simulations. Instead, the DERA*_Gs_*WT structure with only the conf-a of the Lys154 was used as an initial model for both the simulations, as this conformation has been reported in all the previous DERA structures. The S185G and T12I/S185A mutations were computationally introduced in the initial model and the simulations were run using the Charmm36 Force Field for 1 µs each. Three replicas were run for each system. From the simulations, the distribution of the *χ*1b dihedral angle (C-C*_α_*-C*_β_*-C*_γ_*) of K154 showed two predominantly distinct angles for the two variants indicating two conformations of the K154 (Fig 4A). Similar to the crystal structures, the side chain of K154 adopts two distinct conformations in the two variants (Fig 4B). Comparing the structures from the simulations with the crystal structures reveal near-identical conformation of the Lys154 sidechain in the S185G variant (Fig 4B). In the T12I/S185A variant, the Lys154 from the simulations adopts a more exaggerated conformation compared to the crystal structure, but they are both in the same direction, which is distinct and away from the Lys154 conformation in the S185G variant (Fig 4B).

**Figure 4.**
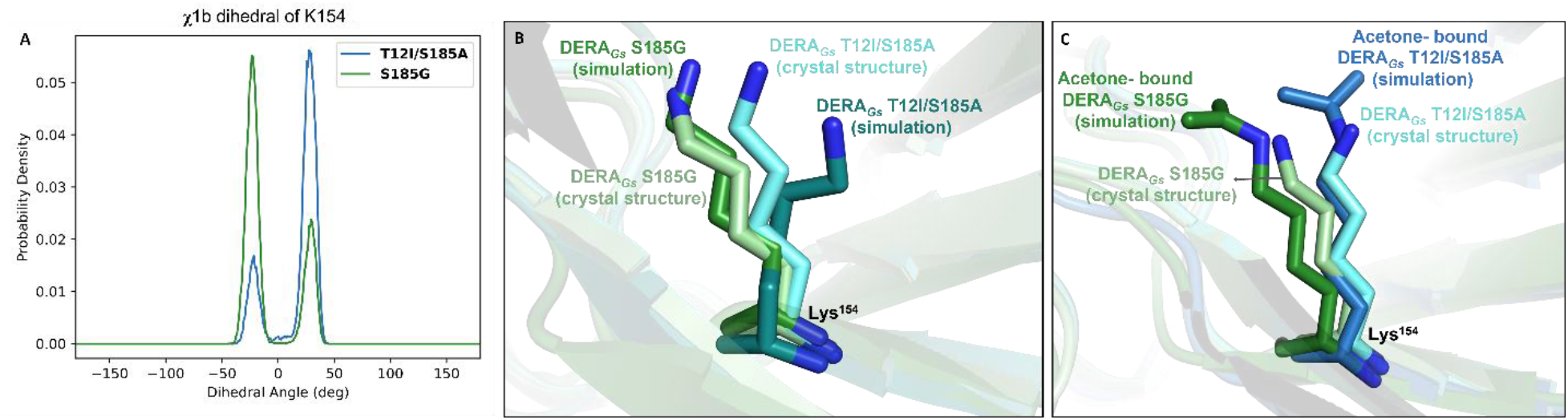
Molecular dynamics simulations reveal two conformations of the catalytic lysine in DERA*_Gs_*, in both the apo-enzyme and the enamine intermediate: (A) Probability density distribution of the χ1b dihedral angle of the catalytic lysine K154 obtained from MD simulations of the DERA*_Gs_* S185G variant and the T12I/S185A variant, showing two distinct populations. (B) Representative structures corresponding to the major conformational states taken from the MD trajectories overlaid with the crystal structures, shows good agreement between the simulated and crystallographic lysine conformations. (C) MD simulation model of the acetone-bound enamine intermediate, generated from the two major conformational states. Simulations of the acetone-bound enamine intermediate also revealed two distinct lysine conformations that are similar to the crystal structures.

After initial attempts to determine the crystal structure of the two DERA*_Gs_* variants with the donor and acceptor substrates, we proceeded with molecular dynamic simulations with the substrate acetone covalently linked to the terminal nitrogen of Lys154 to form the enamine intermediate. The conformations of the Lys154 from the apo-enzyme simulations of the S185G and T12I/S185A variants were used as the initial models for both the variants, and the simulations were run for a total of 3 µs for each system, using the similar approach to the apo-enzyme simulations. The enamine intermediate stabilized into two different conformations of the Lys154 in the S185G and T12I/S185A variants, similar to the apo-enzyme crystal structures (Fig 4C). Taken together, the simulations show that the two different conformations of the Lys154 are regulated by the mutations S185G and T12I/S185A, and that the enamine intermediate also adopts these two conformations.

### Conservation of the enantioselectivity mechanism/regulation in DERA*_Ec_*

DERA*_Ec_* is the industry-relevant homolog of DERA and we wanted to check if the enantioselectivity, via the analogous mutations corresponding to S185G and T12I/S185A from DERA*_Gs_*, is conserved. In DERA*_Ec_*, the position 203 corresponds to Ser185 of DERA*_Gs_* but it is an alanine (Fig. 5A). Due to this alanine, the DERA*_Gs_*variants map onto DERA*_Ec_* with a single substitution each: T18I corresponds to the T12I/S185A double mutant, A203G corresponds to S185G and A203S restores the serine and corresponds to DERA*_Gs_* wild-type. Molecular dynamics simulations using the two variants of DERA*_Ec_*, show two different conformations of the catalytic lysine (K167), consistent with the two conformations of DERA*_Gs_* (Fig 5B, C). Further, the reaction of acetone and thiazole aldehyde using the two DERA*_Ec_* variants resulted in the increased *S*-selectivity (*S/R* = 84:16 e.r.) with the T18I variant compared to A203S (*S/R* = 74:26 e.r.), and the reversal of enantioselectivity to the *R*-isomer in the A203G variant (*S/R* = 24:76 e.r.) (Fig 5D). This is consistent with the DERA*_Gs_* enantioselectivity (Fig 5D). Thus, the conformation of K167 in the T18I variant is close to conf-a resulting in a *S*-selective product, while in the A203G variant, it adopts the novel conf-c resulting in a *R*-selective product. Taken together, we observe that the enantioselectivity is correlated to the conformation of the catalytic lysine residue in DERA enzymes and this is conserved across bacterial species.

**Figure 5.**
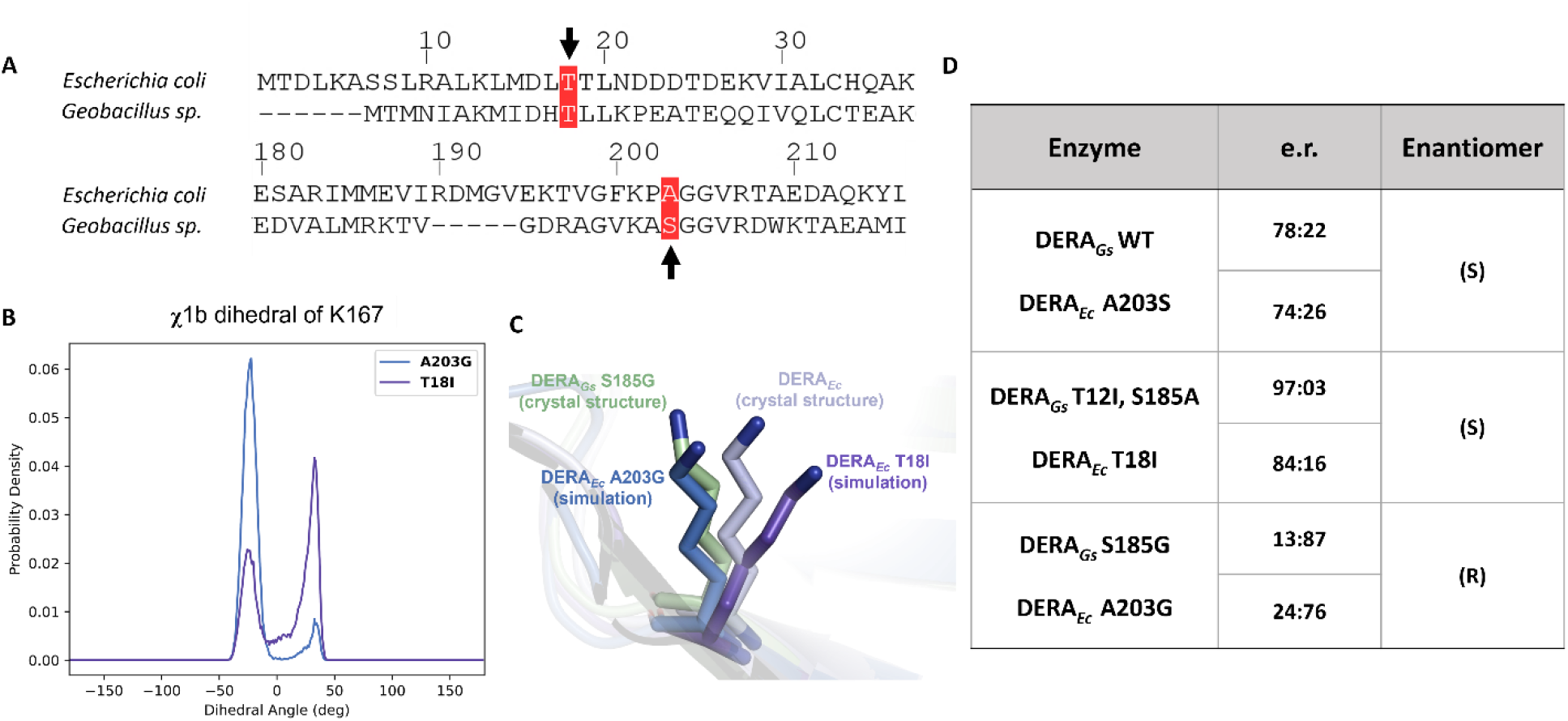
The molecular basis of enantioselectivity in DERA is conserved across bacterial species: (A) Sequence alignment of DERA*_Gs_* and DERA*_Ec_*, with arrows highlighting the residues targeted for mutagenesis. (B) Probability density distributions of the catalytic lysine χ1b dihedral angle obtained from the molecular dynamics simulations of DERA*_Ec_* mutants A203G (corresponding to DERA*_Gs_* S185G) and T18I (corresponding to DERA*_Gs_* T12I/S185A), using the DERA*_Ec_* crystal structure (PDB: 1P1X) as the starting model. The distributions reveal two distinct lysine conformational states. (C) Overlay of representative structures from the MD simulations with the DERA*_Ec_* crystal structure and DERA*_Gs_* S185G variant crystal structure depicts two conformational states of the lysine in DERA*_Ec_* variants. (D) Comparison of the enantioselectivity of DERA*_Gs_* variants with their corresponding DERA*_Ec_* variants shows that equivalent mutations result in similar enantioselectivity, indicating that the molecular basis of enantioselectivity is conserved across bacterial species. The DERA*_Ec_* used in the reactions contains an F200I substitution, which has been shown to improve catalytic efficiency. The mutation is not shown to alter the enantioselectivity of DERA ^26,27^ The sequence alignment is done using Clustal Omega and ESPript.^28^

## Discussion

The DERA enzyme catalyzes the aldol addition reaction and generates a new chiral center in the product^4,29–31^. To understand the mechanism of enantioselectivity, we determined three high-resolution crystal structures of DERA from *Geobacillus sp.* - the wild-type, the T12I/S185A variant and the S185G variant, with the two variants having opposite enantioselectivity. The structures reveal an overall conservation of the protein structure compared to DERA from other organisms and the catalytic pocket is also well conserved. Comparing the three structures of DERA*_Gs_*, reveals no conformational changes in the active site residues, except for the catalytic lysine residue (K154). In the wild-type, the terminal nitrogen of K154 adopts two conformations, we name as conf-a and conf-b; the occupancy of conf-a is 56% and that of conf-b is 44%. This alternate conformation of the catalytic lysine has not been observed before. Given that the DERA*_Gs_* wild-type has two conformations of the K154 and that it results moderate enantioselectivity towards *S*-product, we hypothesize that these two conformations of the catalytic K154 could be a key factor in the enantioselectivity of the enzyme. Consistent with this hypothesis, the K154 adopts different conformations in the two variants of DERA*_Gs_*, which have opposite enantioselectivity. The K154 in the T12I/S185A variant is in conf-a and in the S185G variant it is in conf-c, which is catalytically similar to conf-b. These conformations of the catalytic lysine correlate with their enantioselectivity. The conformations of the catalytic lysine are driven by a combination of steric effect and electrostatic interactions. While the steric effects determine the conformation of the K154 in the T12I/S185A (conf-a) and S185G (conf-c), a combination of steric and electrostatic effects in the wild-type results in two conformations (conf-a, conf-b).

Previous studies of DERA from other organisms report the catalytic lysine in conf-a, but not the conf-b/c. This could be due to two reasons - (i) a lack of sufficient resolution to observe a 1.5 Å shift in the terminal nitrogen atom of the catalytic lysine, or (ii) due to the lack of the citrate molecule (from the crystallization buffer) that could stabilize the conf-b/c. Most of the previously reported DERA structures lack the resolution to observe two conformations of the catalytic lysine (Fig S6). A structure of DERA*_Ec_* is reported at better than 1.0 Å resolution, but the catalytic lysine is observed only in conf-a (Fig S6). This could be because the DERA*_Ec_* active site is subtly different from the DERA*_Gs_*. To ascertain the possible role of the citrate ion, we closely analyzed all the structure of DERA. Among the three structures reported here, the T12I/S185A variant (with conf-a of Lys154) does not have the citrate, while the S185G variant (with conf-c of the Lys154) has citrate in the active site of both the monomers (Fig S2). In the wild type DERA*_Gs_*, the citrate could be modelled only in one monomer due to the poor electron density for the citrate in the second monomer, and the Lys154 adopts both the conf-a and conf-b (Fig S2). A structure of wildtype DERA from *Thermotoga maritima* (PDB code 3R12) also has a citrate ion and the catalytic lysine is modeled in the conf-b (Fig S9).^24^ Taken together, while it is possible that the citrate ion is biasing the K154 towards conf-b/c and these conformations are an artifact of the crystallization conditions, we believe that this is unlikely because (i) the citrate ion forms hydrogen bonding interactions with the sidechain nitrogen of the catalytic lysine K154 in all the conf-a, b and c, and hence, is unlikely to bias the conformation to only conf-b/c (Fig S2), (ii) in the second monomer of the wild-type structure, the citrate is weakly bound, but the K154 still adopts two conformations (Fig S2), (iii) the unbiased molecular dynamic simulations of the two DERA*_Gs_* variants and the corresponding DERA_Ec_ variants, without any citrate ions, show two conformations similar to those in the crystal structures.

In our crystal structures, we have two mutations in the T12I/S185A variant, but the role of the T12I mutation is unclear. Based on our structures, there are no conformational changes at or around the T12I position. In the wild-type and the S185G variant, the oxygen of the T12 sidechain is part of a hydrogen bonding network involving a water molecule, a citrate ion, K183 and the catalytic lysine K154, which is lost in the T12I mutation of the T12I/S185A variant (Fig S7). It is reported that the single T12I mutation results in an increase in the enantioselectivity compared to the S185A variant, but the current structures do not provide any concrete explanation.^9^ It is possible that the T12I affects the orientation of the acceptor substrates or regulates the reactivity of the catalytic lysine, which eventually dictates the subtle change in the stereochemical outcome. Taken together, we conclude that, while the conformation of the catalytic lysine is a key regulator of enantioselectivity, there are other regulators as well.

We propose that the amino acid at the 185 position regulates the conformation of the catalytic lysine and the enantioselectivity of the DERA enzymes. In case of aldol addition of acetone and thiazole aldehyde, the S185A mutation in DERA*_Gs_*results in an enhanced enantioselectivity for *S*-isomer (66% ee) when compared to the wild-type (54% ee)^9^, which is correlated with an increase in the population of conf-a in the observed in the crystal structure of the T12I/S185A variant compared to the wild-type structure. The S185G mutation results in the *R*-isomer with a reversed enantioselectivity (78% ee), which is strongly correlated to the conformational switch to conf-c. Analyzing the sequences of more than 300 DERA proteins, we observe that organisms tolerate either a serine or an alanine in position analogous to 185, but a glycine is never observed (Fig S8). This could be explained using our hypothesis that the glycine at 185 position reverses the stereochemistry of the product, which may not be metabolized by the cell, while the serine or alanine maintain the same stereochemistry that is necessary for the cell.

Our computational and experimental studies with analogous variants of the industry-relevant DERA*_Ec_*, show the critical role of the amino acid at position 203 (185 in DERA*_Gs_*) as a regulator of enantioselectivity and the functional conservation among bacteria. To gain mechanistic insights into the role of the different conformations of the catalytic lysine in determining the stereochemical outcome, we ran molecular dynamic simulations with the enamine intermediate, which resulted in two different conformations for the enantiocomplementary variants. Mechanistically, since both the reaction substrates are achiral, the stereochemical outcome is dictated entirely by the environment of the enzyme active site during catalysis.^21^ The absolute configuration of the product is most likely determined by the orientation of the enamine intermediate with respect to the acceptor aldehyde, and by its attack on one enantiotopic face of the prochiral acceptor aldehyde.^31–33^ Our data shows that the catalytic lysine can adopt two conformations that dictate the conformation of the enamine intermediate, which could react with either of the faces of the acceptor substrate, determining the ultimate stereochemistry of the product (Fig 6). This mechanism of enantioselectivity has to be confirmed with further studies and the possible conservation of this mechanism among the TIM-barrel family of enzymes with achiral substrates needs to be probed.

**Figure 6.**
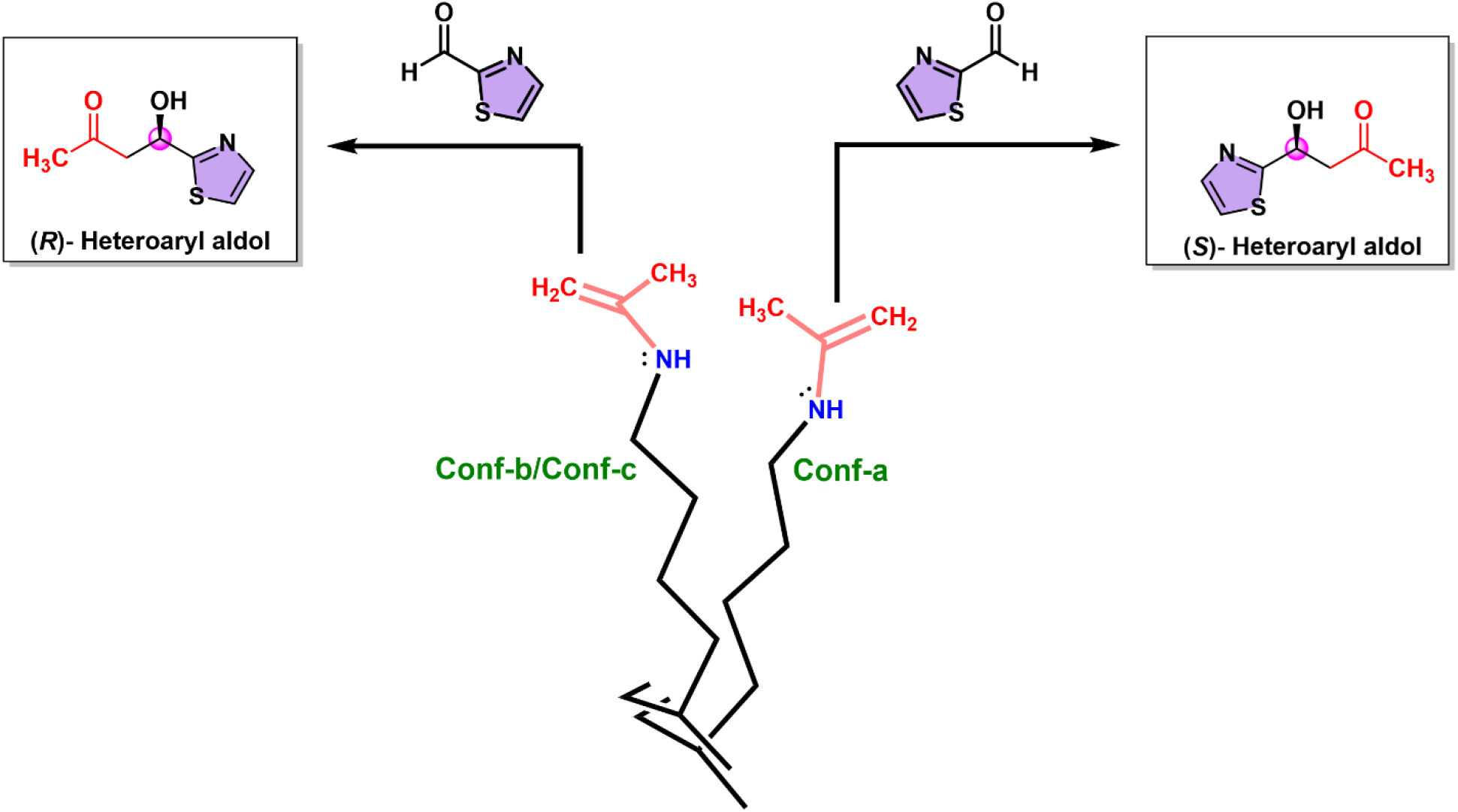
The proposed model for the enantioselectivity of DERA*_Gs_*. The alternative conformations of the Lys154 residue can generate enamine intermediates, enabling nucleophilic attack on either face of the incoming pro-chiral electrophile giving rise to an enantiomeric mixture of aldol products in the wild-type DERA enzymes. In case of variants that can stabilize one of these conformations, the product would be predominantly either (*S*)- or (*R*)-aldol.

## Acknowledgements

We acknowledge support of the Department of Atomic Energy, Government of India, under Project Identification No. RTI 4007. The authors acknowledge Elettra Sincrotrone Trieste for providing access to the X-ray diffraction beamline and thank Raghurama P Hegde for his guidance during data collection. TS gratefully acknowledges financial support from the Science and Engineering Research Board, Department of Science and Technology, Government of India (SERB-DST, now ANRF; Grant No. SRG/2021/001015); the Scheme for Transformational and Advanced Research in Sciences, Ministry of Education, Government of India (MoE-STARS; Grant No. MoE-STARS/STARS-2/2023-0992); and the Anusandhan National Research Foundation, Ministry of Science and Technology, Government of India (ANRF), through the Partnerships for Accelerated Innovation and Research (PAIR; Grant No. ANRF/PAIR/2025/000012/EPAIR, Task Nos. 64 and 74) and the Advanced Research Grant (ARG; Grant No. ANRF/ARG/2025/009779/CS). We would like to thank Deepa S, Kaustubh R. Mote, Sreejith Raran Kurussi, Kiran Kulkarni and Aravind Penmatsa for their inputs and contributions.

## Author contributions

TS & ATV conceived the project, SD & ATV determined the structures, KJ & TS performed the enzymatic reactions, AN & JM ran the molecular dynamic simulations. All the authors analyzed the data. SD & ATV wrote the manuscript and all the authors provided the inputs. All the authors approved the final version of the manuscript.

