## Supplementary Data for "A Conformational Switch in the Catalytic Lysine Governs the Stereochemical Outcome of DERA-Catalyzed Aldol Addition"

### Supplementary Figures

**Table S1**. Data Collection and Refinement Statistics.

| | $DERA_{Gs}$ Wild type<br>PDB ID 44HZ | $DERA_{Gs}$ –S185G<br>PDB ID 44FM | $DERA_{Gs}$ –T12I/S185A<br>PDB ID 44IA |
|---|---|---|---|
| **Data collection** | | | |
| Space group | P 1 21 1 | P 1 21 1 | P 1 21 1 |
| Cell dimensions a, b, c (Å) | 45.33, 70.57, 83.68 | 45.29, 67.98, 59.90 | 45.37, 70.20, 60.34 |
| α, β, γ (°) | 90.00, 94.08, 90.00 | 90.00, 94.66, 90.00 | 90.00, 94.11, 90.00 |
| Wavelength (Å) | 1 | 1 | 1 |
| Resolution range (Å) | 41.75 - 1.19 (1.21 – 1.19) | 37.605 – 1.50 (1.53 – 1.50) | 38.03 – 1.22 (1.24 – 1.22) |
| Unique reflections | 167749 (8317) | 57613 (2814) | 108864 (5230) |
| Multiplicity | 6.2 (5.5) | 6.1 (6.0) | 6.1 (6.0) |
| Completeness (%) | 99.7 (99.8) | 99.4 (99.1) | 97.4 (95.4) |
| Mean I/σ(I) | 15.5 (3.0) | 11.2 (3.0) | 13.0 (2.9) |
| CC1/2 | 0.999 (0.782) | 0.996 (0.849) | 0.997 (0.855) |
| Rmerge | 0.068 (0.646) | 0.116 (0.789) | 0.079 (0.478) |
| Rpim | 0.030 (0.300) | 0.051 (0.359) | 0.035 (0.212) |
| Rmeas | 0.074 (0.714) | 0.127 (0.869) | 0.087 (0.524) |
| Wilson B factor ($Å^2$) | 12.1 | 7.2 | 10.1 |
| **Refinement** | | | |
| Resolution (Å) | 1.19 | 1.50 | 1.22 |
| Rwork/Rfree (%) | 12.4/14.0 | 16.3/18.6 | 18.6/20.4 |
| No. atoms: Protein | 6315 | 6329 | 6304 |
| Ligand/Ion | 18 | 36 | 0 |
| Water | 341 | 179 | 190 |
| RMSD bond lengths (Å) | 0.0127 | 0.0087 | 0.0106 |
| RMSD bond angles (°) | 1.996 | 1.696 | 1.801 |
| Ramachandran favored (%) | 98 | 99 | 98 |
| Ramachandran allowed (%) | 2 | 1 | 2 |
| Ramachandran outliers (%) | 0 | 0 | 0 |
| Rotamer outliers (%) | 1 | 1 | 0 |
| Clashscore | 4 | 1 | 2 |

Values in parentheses refer to the highest-resolution shell. Rwork and Rfree were calculated using the working and test reflection sets, respectively. The free-reflection set was used for cross-validation during refinement.

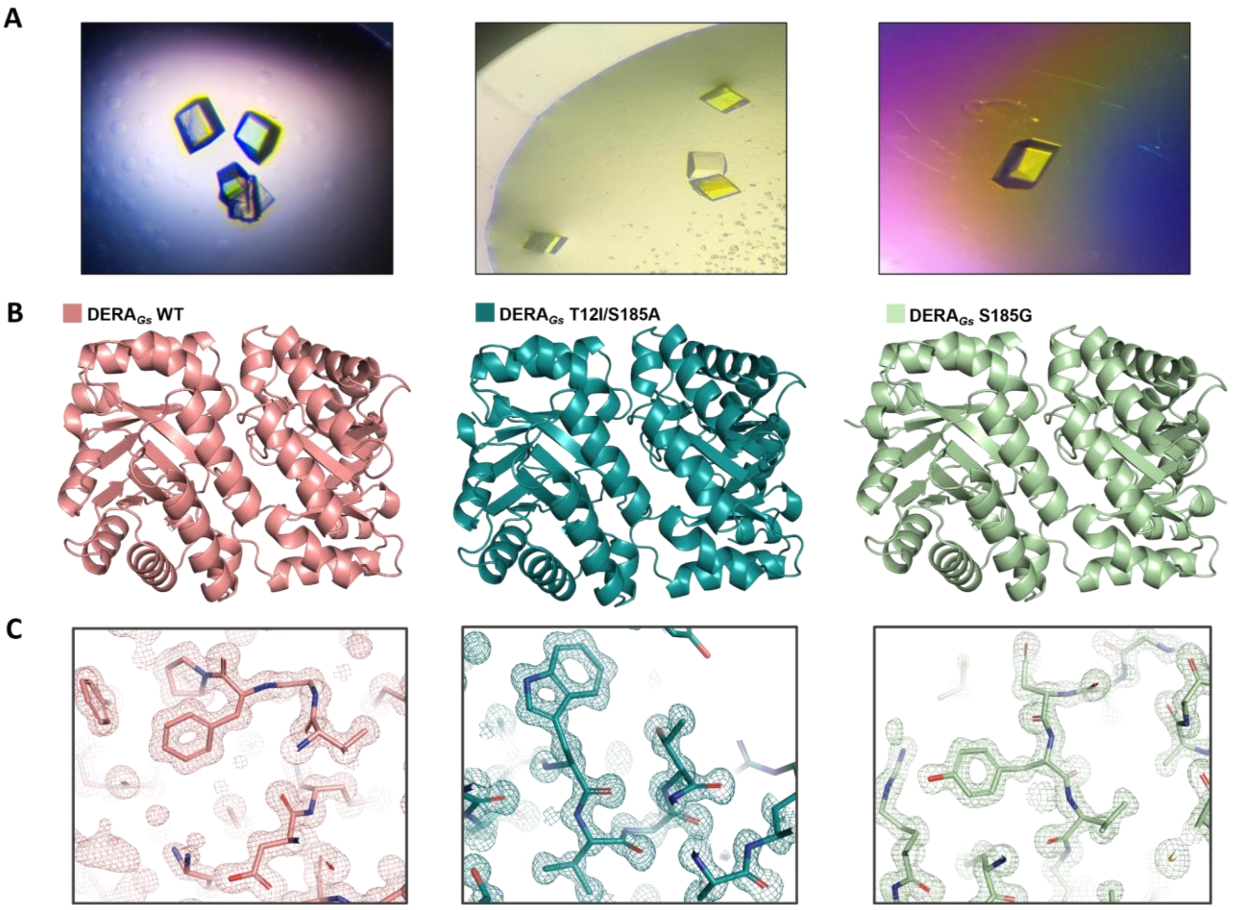


**Figure S1. Crystal structures of DERA$_{Gs}$ protein and electron density.** (A) Images of the cuboidal crystals of wild-type DERA$_{Gs}$, T12I/S185A, and S185G proteins, with dimensions of ~200 µm. (B) Cartoon representations of the dimers of DERA$_{Gs}$ wild-type (WT) (salmon), T12I/S185A (cyan) and S185G (green) proteins. (C) Below each dimer a corresponding region of the 2F$_o$-F$_c$ electron density map, contoured at 1.5σ, is shown with the fitted atomic model.

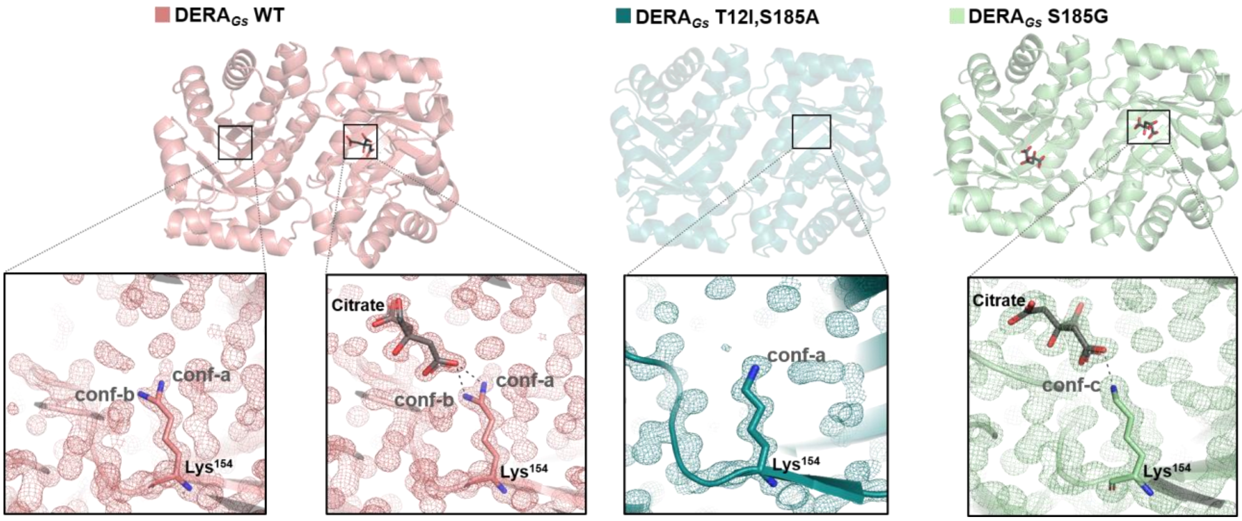


**Figure S2. Presence of a citrate ion in the WT and DERA variants.** Overall structures of WT DERA$_{Gs}$ (salmon), the T12I/S185A variant (cyan), and the S185G variant (pale green) are shown in cartoon representation. Insets highlight the active-site region with the corresponding 2$F_o$-$F_c$ electron density maps contoured at 1.5σ. A citrate ion, from the crystallization condition, could be modelled in only one monomer of the WT dimer due to poorly defined electron density in the second monomer. No citrate ion is present in the T12I/S185A variant, whereas citrate was observed in both monomers of the S185G variant. Lys154 and the citrate are shown in stick representation.

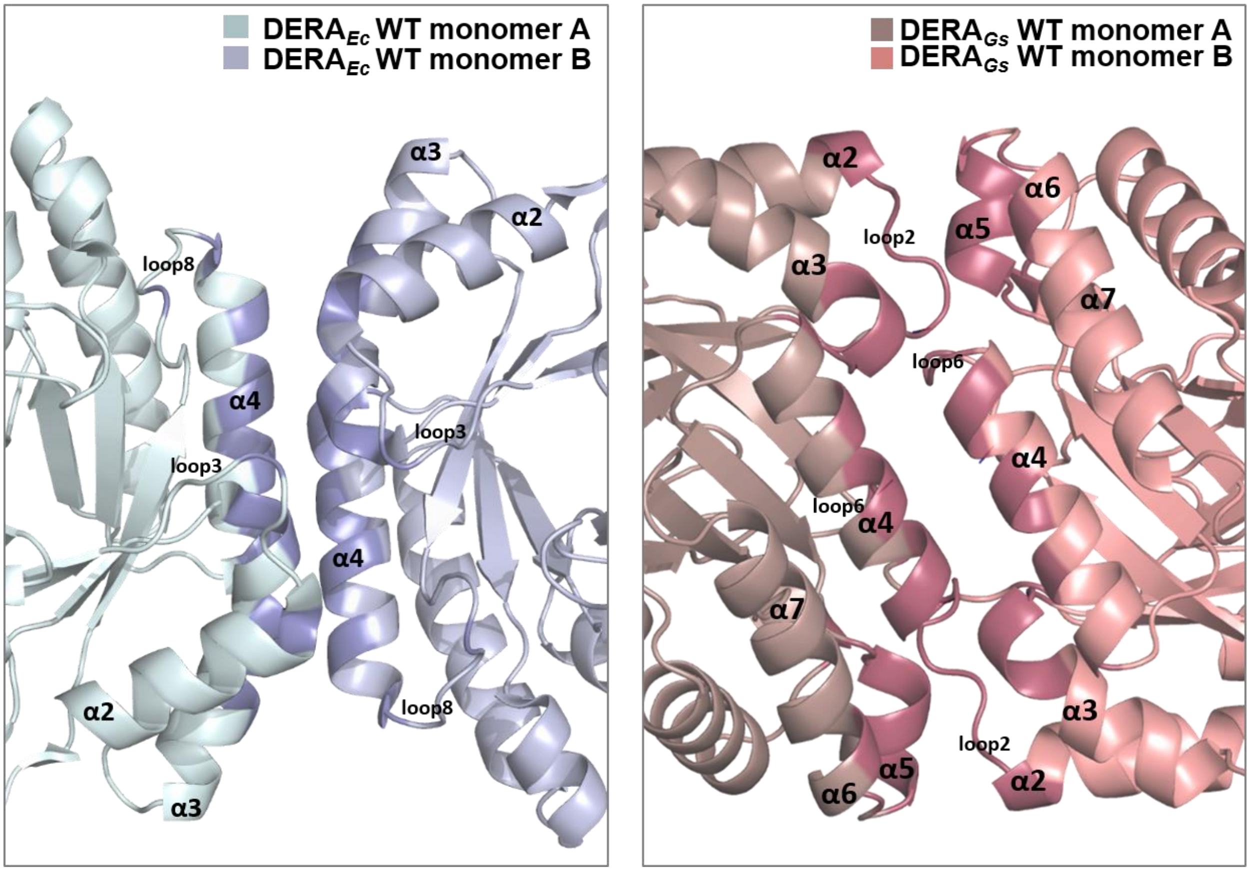


**Figure S3. Comparison of dimer interfaces in DERA*Ec* and DERA*Gs*.** Cartoon representations of the dimers of DERA*Ec* (left) and DERA*Gs* (right) highlighting residues that contribute to the dimer interface (shown as sticks). The DERA*Gs* dimer exhibits a larger interface, involving helices α2–α7 and loops 2, 4, 6, 8, 9, and 14 from both monomers. In contrast, the DERA*Ec* dimer interface is more restricted, primarily comprising helices α2–α4 and loops 3 and 8. The solvent buried surface area of DERA*Ec* is 595Å$^2$ and that of DERA*Gs* is 1277Å$^2$. These surface areas were calculated using PDBePISA v1.52 (Krissinel and Henrick, 2007)

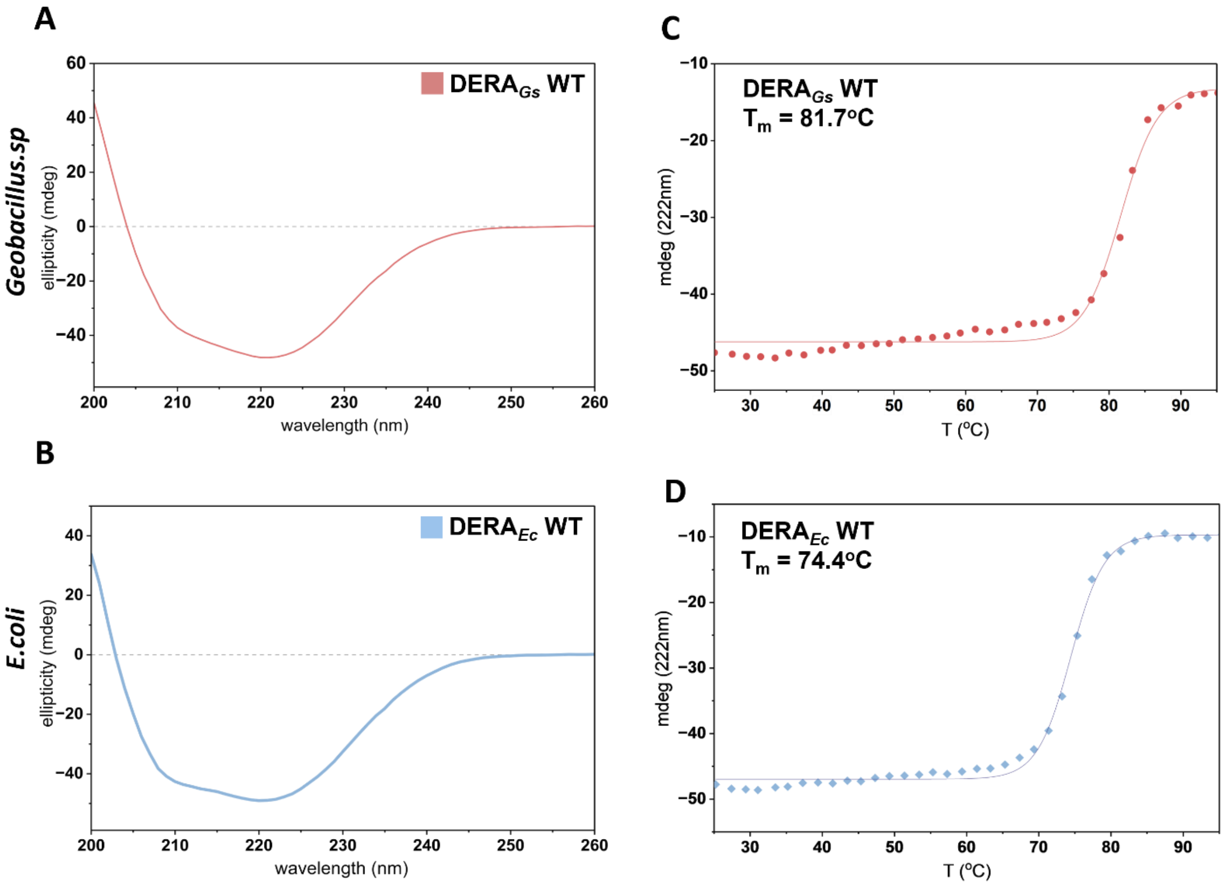


**Figure S4: Determination of melting point of DERA$_{Gs}$ and DERA$_{Ec}$.** (A, B) Far-UV CD spectra of DERA$_{Gs}$ and DERA$_{Ec}$, respectively, showing a minimum at 222 nm characteristic of α-helical secondary structure. (C, D) Thermal unfolding curves of DERA$_{Gs}$ and DERA$_{Ec}$, monitored at 222 nm, and the associated melting temperature (Tm).

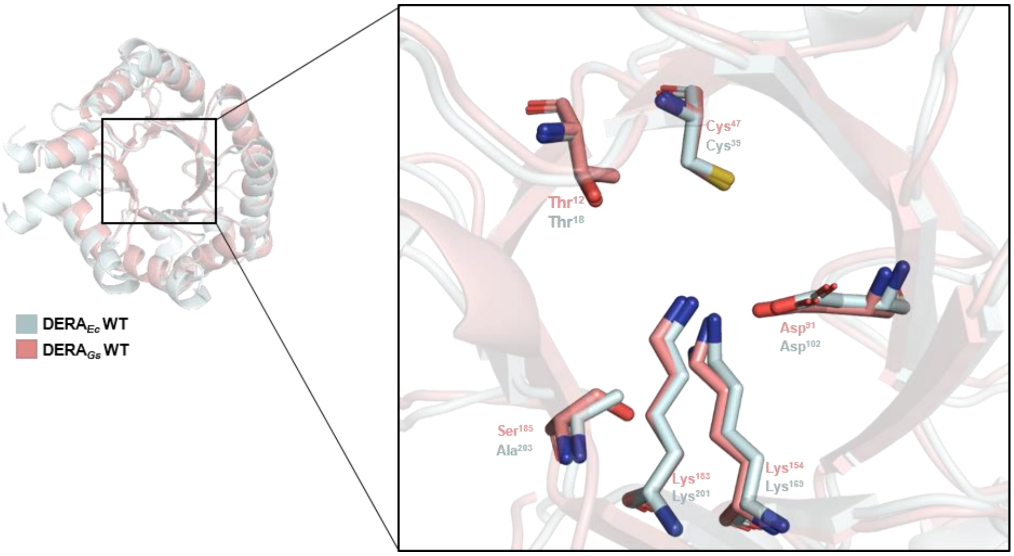


**Figure S5. Enlarged view of the active sites of DERA$_{Gs}$ WT and DERA$_{Ec}$ WT (PDB 1P1X):** The conserved catalytic residues superimpose very well despite a low sequence identity of 34.8%.

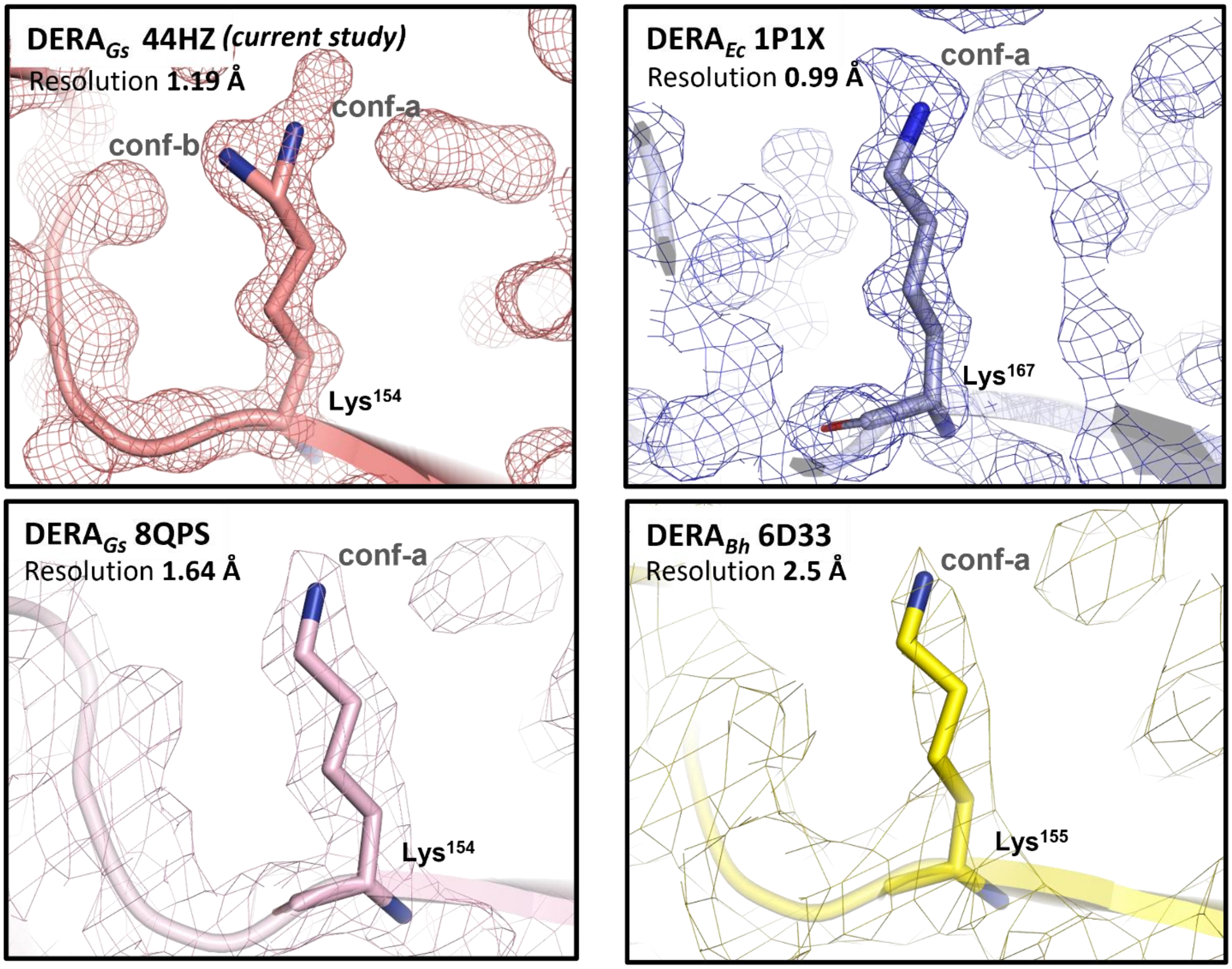


**Figure S6. Conf-b is a novel conformation of the catalytic lysine in DERA:** The catalytic lysine of DERA from various organisms, along with the $2F_o - F_c$ electron-density map and the corresponding resolutions are shown. The $DERA_{Gs}$ structure determined in this study (top left) exhibits well-defined electron density that supports two distinct conformations of the catalytic Lys154 side chain. In contrast, the catalytic lysine in the previously reported structures of *E. coli* ($DERA_{Ec}$) and *Bacillus halodurans* ($DERA_{Bh}$), a single conformation (conf-a) is observed. All $2F_o - F_c$ electron-density maps are contoured at 1.0σ.

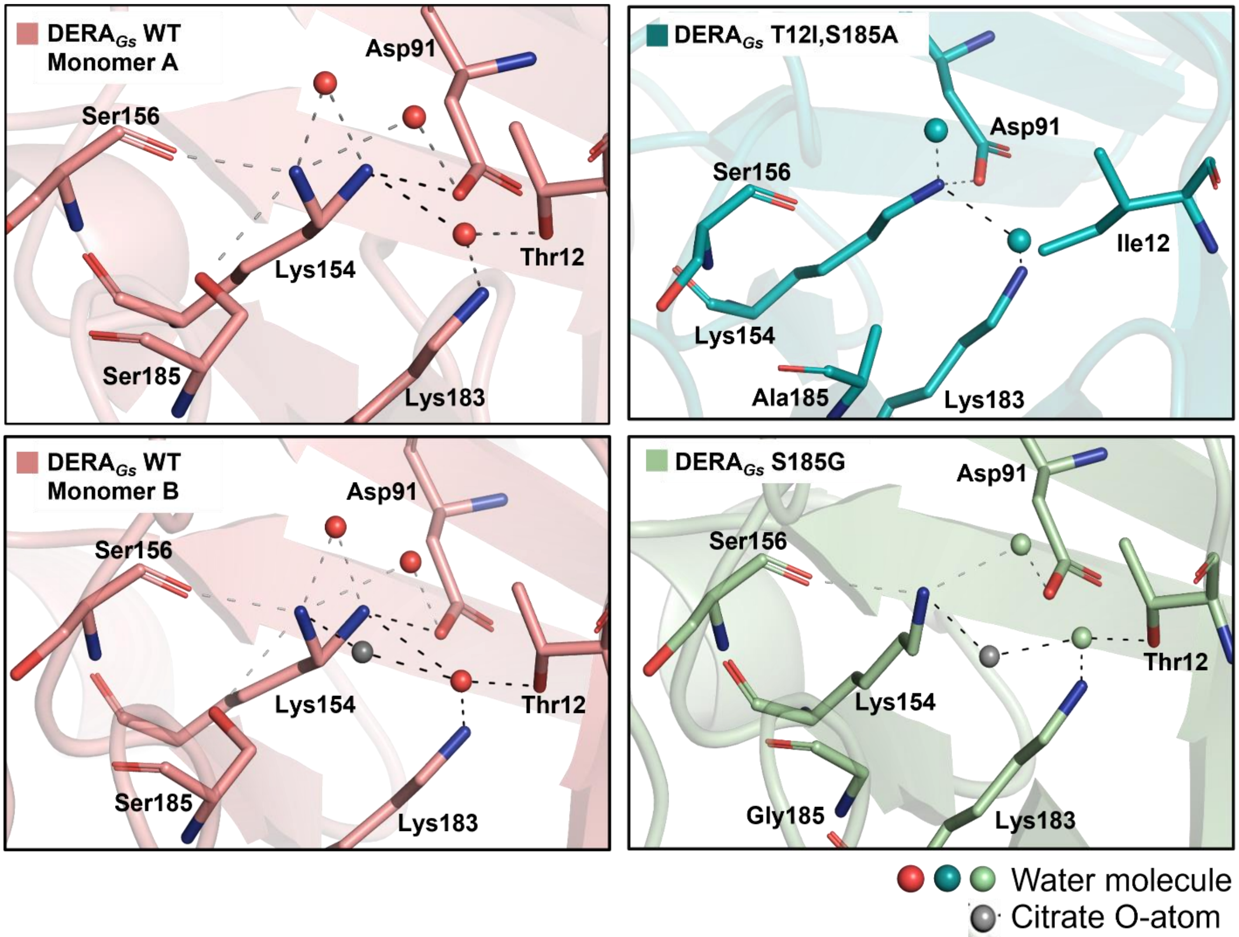


**Figure S7. Stabilizing interactions of the different conformations of the catalytic lysine:** The active-site solvent network stabilizes both conformations of the catalytic Lys154 residue in DERA$_{Gs}$ WT and all mutant structures. In the T12I/S185A variant, the T12I substitution results in the loss of hydrogen-bonding interactions with one conserved water molecule that mediate interactions between Lys154 and Lys183, thereby altering the solvent network around the active site. The citrate molecules, shown as a grey ball, are occupying the site of the molecule's terminal carboxyl oxygen atom. It is modelled in one monomer of DERA$_{Gs}$ WT, both monomers of S185G mutant, and is absent from both monomers of the T12I/S185A mutant.

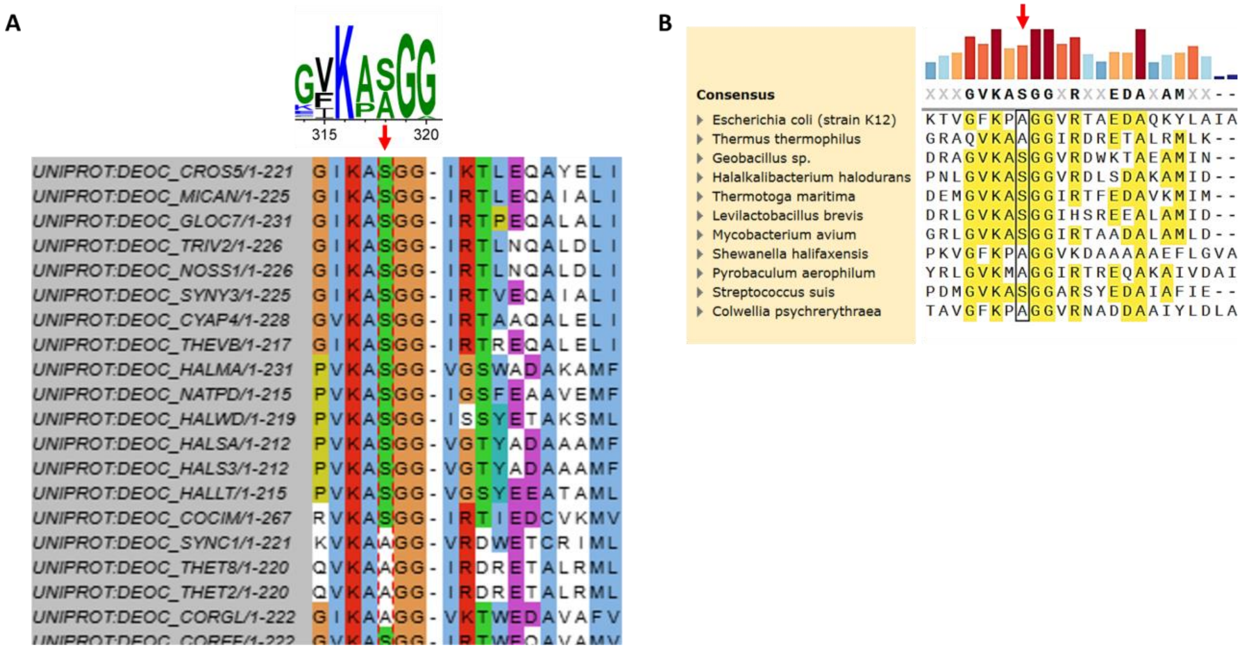


**Figure S8. Conservation of the 185th residue in the active-site position across DERA homologues (**A) Multiple sequence alignment of 340 DERA homologues identified by PSI-BLAST, highlighting the residues corresponding to Ser185 in DERA$_{Gs}$. The sequences were aligned using Clustal Omega. All sequences were visualised using JalviewJS, out of which 20 have been shown here. Sequence conservation is summarized by the WebLogo, which shows that this position (indicated with red arrow) is occupied almost always by either serine or alanine. The conservation of these residues and the lack of glycine indicates that the alternative Lys154 conf-c conformation is not observed in naturally occurring DERA homologues resulting in only one enantiomer, which corroborates our hypothesis. (B) Multiple sequence alignment of all available crystal structures of DERAs available in the Protein Data Bank (PDB) also shows the conservation of either serine or alanine at the same position. The sequence alignments were generated in SnapGene using Clustal Omega.

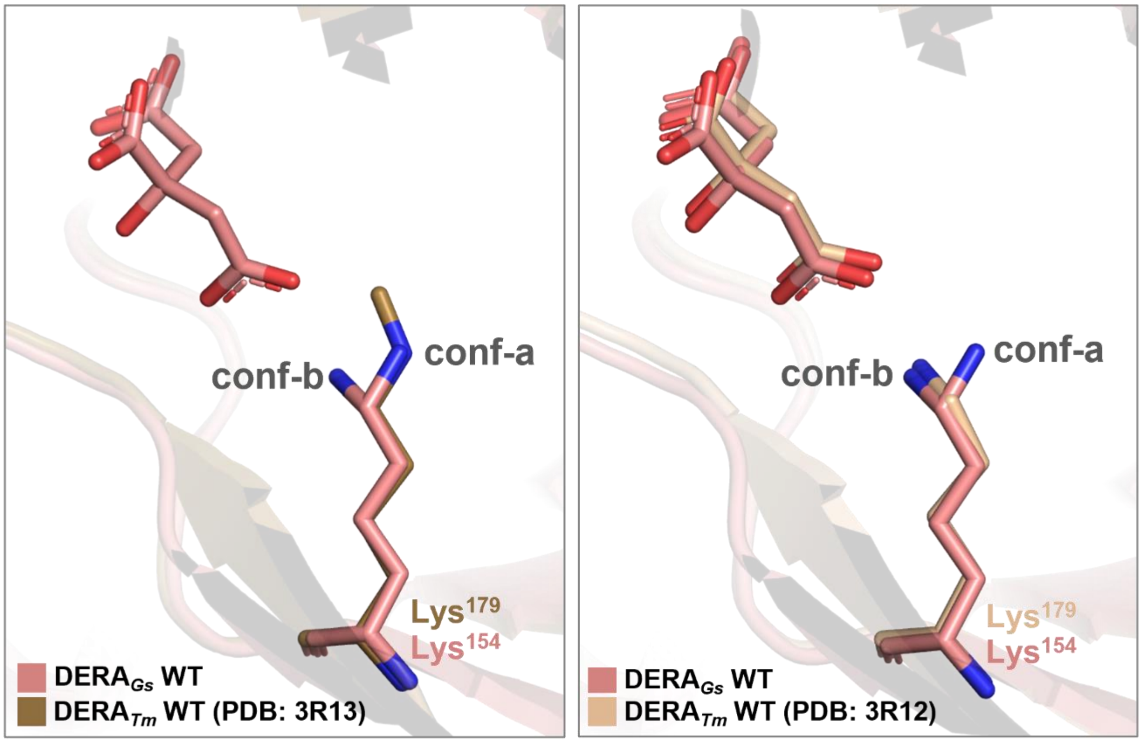


**Figure S9: Structural alignment of *Thermotoga maritima* DERA (DERA$_{Tm}$) with DERA$_{Gs}$.** (A) The crystal structure of DERA$_{Tm}$ WT, without the citrate ion (PDB 3R13), is aligned with DERA$_{Gs}$ WT, and the catalytic Lys179 (corresponding to Lys154 in DERA$_{Gs}$) adopts the conf-a. (B) The crystal structure of DERA$_{Tm}$ WT, with a citrate ion (PDB 3R12), is aligned with DERA$_{Gs}$ WT, and the catalytic Lys179 (corresponding to Lys154 in DERA$_{Gs}$) adopts the conf-b.

#### Methods

##### Cloning and purification

Chemically competent *E. coli* BL21 (DE3) cells were transformed with the plasmid encoding the C-terminal His-tagged DERA variant. A single transformant colony was inoculated into 10 mL of LB medium supplemented with 30 µg/mL ampicillin and grown at 37 °C with shaking at 200 rpm for 12–16 h. Subsequently, 1 mL of the overnight culture was transferred into 1 L of TB medium containing 30 µg/mL ampicillin. The culture was incubated at 37 °C and 200 rpm until an $OD_{600}$ of approximately 0.8 was reached (3 h). Protein expression was then induced by adding 500 µM isopropyl-β-D-thiogalactopyranoside (IPTG), followed by overnight incubation at 25 °C with shaking at 170 rpm. The cells were harvested by centrifugation and the resulting pellet was resuspended in 20 mM potassium phosphate buffer (pH 7). Cells were lysed by ultrasonication, after which the lysate was centrifuged to separate the soluble fraction. The clarified lysate was filtered using a 0.45 µm filter and loaded on a 5 mL HisTrap FF column. After washing the column with 100 mL of 20 mM potassium phosphate containing 20 mM imidazole (pH 7), the bound protein was eluted using 7 mL of 20 mM potassium phosphate containing 300 mM imidazole (pH 7). The eluted protein was subsequently exchanged into 20 mM potassium phosphate buffer (pH 7) by size-exclusion chromatography using a HiLoad 26/600 Superdex 200 prep grade column (Cytiva). Protein concentration was estimated from the absorbance at 280nm. The oligomeric state and molecular mass of the purified protein were verified by SEC-MALS. Finally, the remaining purified protein was snap-frozen in liquid nitrogen and stored at −20 °C until further use.

##### X-ray crystallography

Crystallization trials were initiated using the sitting-drop vapour-diffusion method in MRC 96-well 3-drop crystallization plates (Molecular Dimensions) at 25 °C. Initial screening employed the JCSG-plus (Molecular Dimensions) and Crystal Screen 1 & 2 (Hampton Research) crystallization kits. 150 nL drops were dispensed using a Mosquito liquid-handling system (SPT Labtech) by combining purified $DERA_{Gs}$ at concentrations of 5 mg/mL, 10 mg/mL, or 20 mg/mL in 20 mM potassium phosphate (pH 7) with reservoir solution in a 1:1 volume ratio. Among the screened conditions, the most promising crystals were obtained from condition

C6 of the JCSG-plus screen, consisting of 0.1 M phosphate-citrate (pH 4.2) and 40% PEG 300. Crystal quality was improved by varying the pH and PEG 300 concentration. Diffraction-quality crystals appeared within a few weeks in solutions containing 0.1 M phosphate/citrate (pH 4.4–4.6) and 42–50% PEG 300. Prior to data collection, crystals were transferred to reservoir solution supplemented with 25% (v/v) glycerol for cryoprotection and flash-cooled in liquid $N_2$. X-ray diffraction data were collected at the XRD2 beamline, Elettra Sincrotrone Trieste[1] (2.4 GeV synchrotron), Trieste, Italy, and processed using XDSAPP3.[2,3] The crystals belonged to space group $P2_1$, contained two polypeptide chains in the asymmetric unit. The structure of apo DERA$_{Gs}$, determined to a resolution of 1.19 Å, was solved by molecular replacement with MOLREP,[4,5] using the crystal structure of DERA from *Bacillus halodurans* (PDB entry 6D33)[6] as the search model. Iterative cycles of manual model rebuilding in WinCOOT[7] and refinement with REFMAC5[8] were carried out. The final refinement stages included the addition and validation of water molecules and citrate molecules, followed by anisotropic atomic B-factor refinement. The DERA$_{Gs}$ S185G and DERA$_{Gs}$ T12I, S185A mutant structures were refined at resolutions of 1.50 Å and 1.22 Å, respectively, using REFMAC5 and following the same refinement strategy applied to the apo DERA$_{Gs}$ structure. The final structural models were assessed using the wwPDB Validation Server[9] (https://validate.wwpdb.org). Structural analysis was performed in WinCOOT, and publication-quality Figures were prepared using PyMOL (Schrödinger). Atomic coordinates and structure factors were deposited in the Protein Data Bank under accession codes. A summary of the crystallographic data collection and refinement statistics is provided in.

**Biochemical characterisation**

Circular dichroism (CD) spectroscopy was used to determine the melting temperature of the proteins. The CD signal at 222 nm was monitored as the proteins were heated from 25 °C to 95 °C at a rate of 2 °C/min. The change in molar ellipticity with temperature produced a sigmoidal curve. The inflection point of this curve was taken as the melting temperature ($T_m$) of the protein.

**Simulation**

The starting structure for wild-type configuration was taken from the experimentally determined crystal structure of the wild-type DERA$_{Gs}$ (PDB 44HZ) with the K154 conf-a. We generated the structures of T12I/S185A and S185G by mutating T12 to I12 and S185 to A185 for Double Mutant and S185 to G185 using CHARMM GUI PDB reader and manipulator.[10] All three systems were solvated using solution builder[11,12] and Charmm TIP3P[13] water. The salt concentration was 0.15 M KCl and pH 7. We used the CHARMM36[14] force field for all of the systems. We did a position restrained NVT equilibration for 5ns; at an average temperature of 298.15K using the V-rescale Thermostat, with relaxation time of 1.0 ps.[15] The same temperature was kept throughout each of the simulations using the above-mentioned thermostats. Then we did a restrain free NPT simulation for 5 ns. For the NPT simulation, we used the C-rescale barostat[16] to keep an average pressure of 1.0bar constant. The time constant value of 5.0 ps was used with compressibility factor 4.5e-5 (1/bar). Restraint free simulations were performed on the equilibrated structures. Three replicas of each system were simulated for 1000 ns long using GPU-enabled Gromacs 20xx package.[17] And these simulations were concatenated to give 3 microsecond long trajectories for each, DERA$_{Gs}$ WT, T12I/S185A and S185G systems. To create the intermediate, we prepared acetylene bound lysine using Charmm GUI ligand designer.[18,19]

Using the same protocol we ran simulations for DERA$_{Ec}$ A203G; A203S; T18I and wild-type variants, starting from the PDB structure 1P1X.[20] All the analyses were performed using the MDAnalysis module.[21] The dihedrals were calculated using the gmx angle module. The visualization of trajectories was done using VMD.[22]

We took snapshots from the mutated DERA$_{Gs}$ T12I/S185A and S185G variants corresponding to their highest populated conformations of Lys154. The systems were built and solvated in Gromacs. Ions were used in neutralising concentration. The enamine intermediate system was generated for all three: DERA$_{Gs}$ wild-type, S185G and T12I/S185A systems. Each system was equilibrated using the Nose-Hoover Thermostat, with relaxation time of 1.0 ps.[23,24] Then we did a restrain free NVT equilibration for 5 ns followed by a restrained free NPT simulation for 5 ns. For the NPT simulation, we used the Parrinello-Rahman barostat.[25]

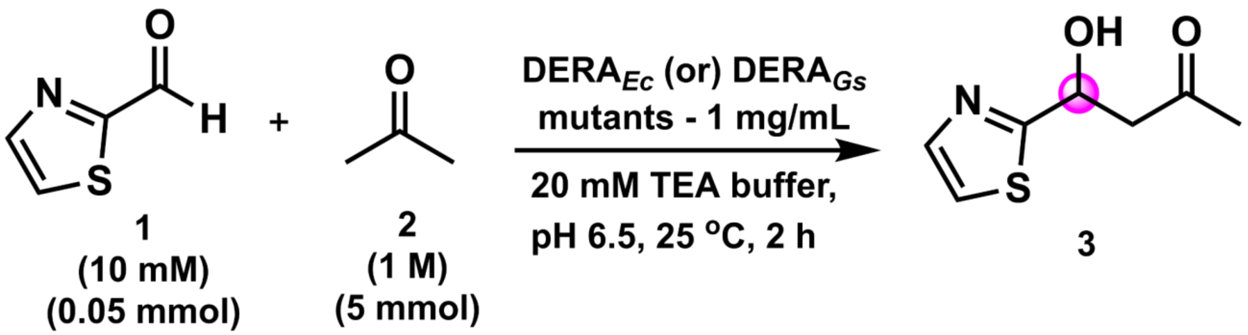


**Materials:** The aldehydes were purchased from BLD-Pharm and acetone from FINAR. Reactions were monitored using thin-layer chromatography (TLC) with aluminium sheets of silica gel 60 F254 from Merck. TLC plates were visualized with UV light (254 nm) or *p*-anisaldehyde stain.

**General Procedure for Analytical-Scale Reactions:**

The reactions were performed on a 10 mM (0.05mmol) scale with a total reaction volume of 5 mL. In a 25-mL conical flask, 20 mM of TEA buffer, pH 6.5, was taken. Aldehyde (10 mM final concentration) solution in DMSO (5% v/v) was added to the buffer, followed by acetone (1 M final concentration) and purified enzyme (1 mg/mL; 0.08 mol%). The reaction mixture was incubated in an orbital shaker at 25 $^{o}$C, 120 rpm for 2 hours. Upon completion, the reaction mixture was extracted using ethyl acetate (10 mL * 3). The organic layer was dried over sodium sulphate, filtered, and the solvent was evaporated using a rotary evaporator. The crude mixture was analyzed by $^{1}$H NMR and the spectral values were matching with previously reported data.[26] The enantiomeric ratio was determined by chiral HPLC analysis using Diacel Chiralpak AD-H column, hexane/isopropanol 95:05, flow rate 1 mL min$^{-1}$, $\lambda$ = 254 nm.

<table>
<tr><th>Enzyme</th><th>e.r.</th><th>Enantiomer</th></tr>
<tr><td rowspan="2">DERAGs<br>DERAEc A203S</td><td>78:22</td><td rowspan="2">(S)</td></tr>
<tr><td>74:26</td></tr>
<tr><td rowspan="2">DERAGs T12I, S185A<br>DERAEc T18I</td><td>97:03</td><td rowspan="2">(S)</td></tr>
<tr><td>84:16</td></tr>
<tr><td rowspan="2">DERAGs S185G<br>DERAEc A203G</td><td>13:87</td><td rowspan="2">(R)</td></tr>
<tr><td>24:76</td></tr>
<tr><td>DERAEc</td><td>84:16</td><td>(S)</td></tr>
</table>

Note: The $DERA_{Ec}$ used in the reactions contains an F200I substitution, which has been shown to improve catalytic efficiency.[27,28]

**HPLC Profiles:**

Racemic standard

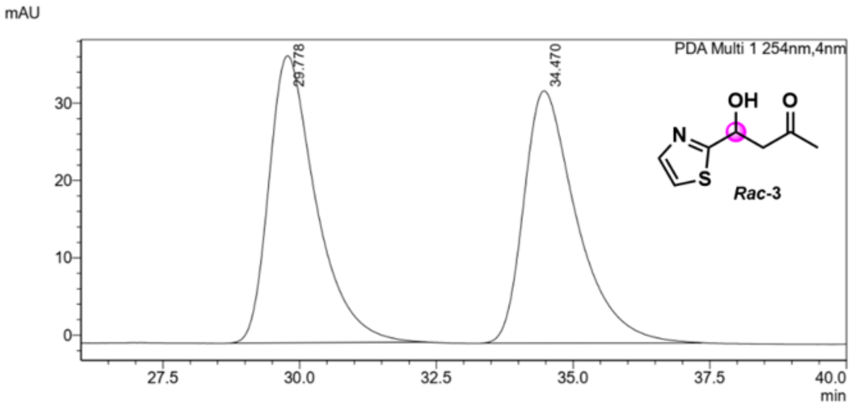


<Peak Table>

PDA Ch1 254nm

| Peak# | Ret. Time | Area | Area% |
|---|---|---|---|
| 1 | 29.778 | 2223273 | 50.300 |
| 2 | 34.470 | 2196737 | 49.700 |
| Total | | 4420010 | 100.000 |

$DERA_{Gs}$

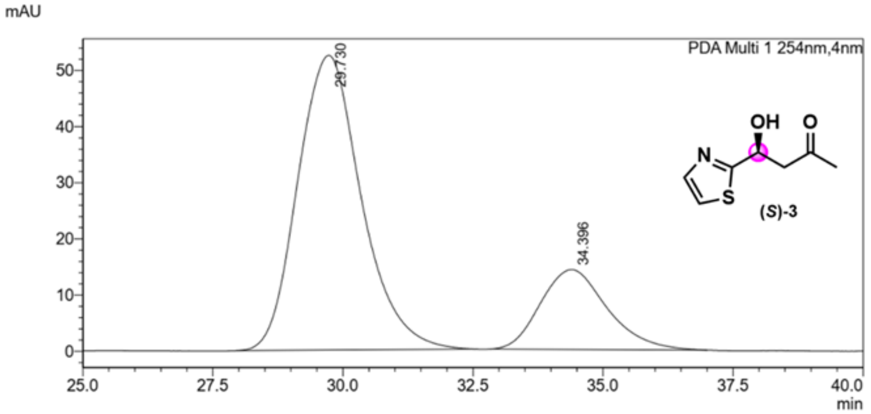


<Peak Table>

PDA Ch1 254nm

| Peak# | Ret. Time | Area | Area% |
|---|---|---|---|
| 1 | 29.730 | 4471260 | 77.873 |
| 2 | 34.396 | 1270446 | 22.127 |
| Total | | 5741706 | 100.000 |

$DERA_{Ec}$-A203S

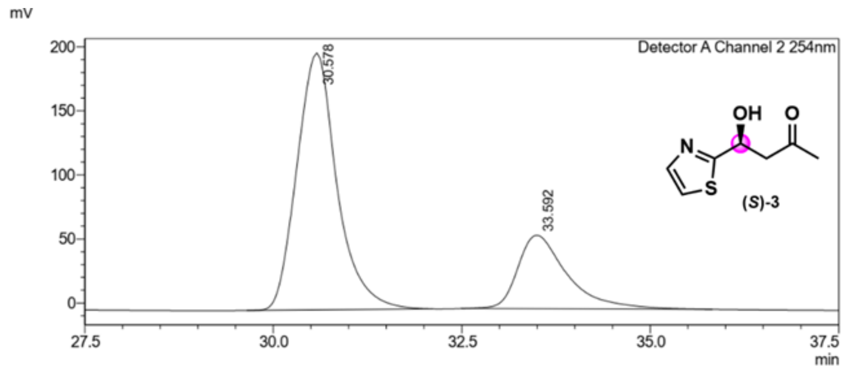


Detector A Channel 2 254nm

| Peak# | Ret. Time | Area | Area% |
|---|---|---|---|
| 1 | 30.578 | 7414335 | 73.778 |
| 2 | 33.592 | 2635160 | 26.222 |
| Total | | 10049495 | 100.000 |

#### DERA$_{Gs}$-T12I/S185A

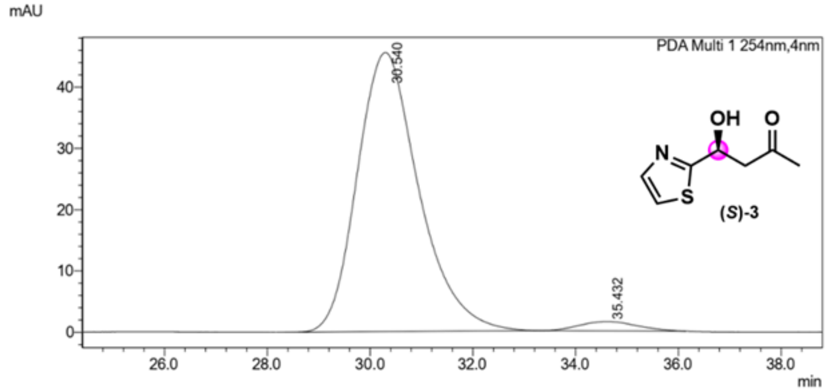


**<Peak Table>**

PDA Ch1 254nm

| Peak# | Ret. Time | Area | Area% |
|---|---|---|---|
| 1 | 30.540 | 4730959 | 96.959 |
| 2 | 35.432 | 148405 | 3.041 |
| Total | | 4879364 | 100.000 |

#### DERA$_{Ec}$-T18I

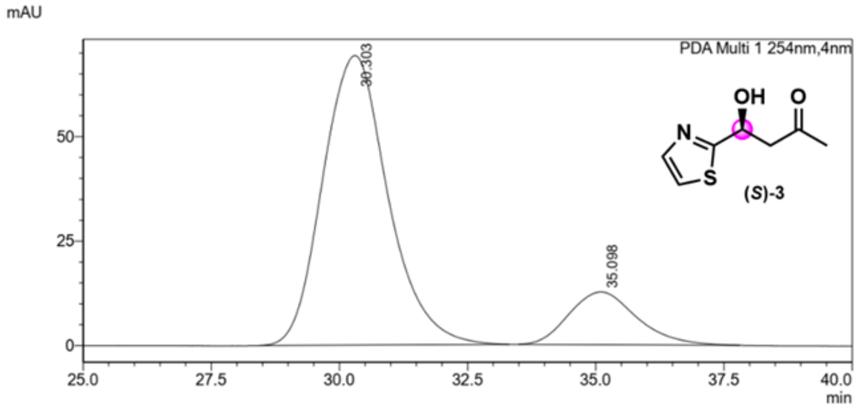


**<Peak Table>**

PDA Ch1 254nm

| Peak# | Ret. Time | Area | Area% |
|---|---|---|---|
| 1 | 30.303 | 6144096 | 84.072 |
| 2 | 35.098 | 1163998 | 15.928 |
| Total | | 7308094 | 100.000 |

DERA$_{Gs}$-S185G

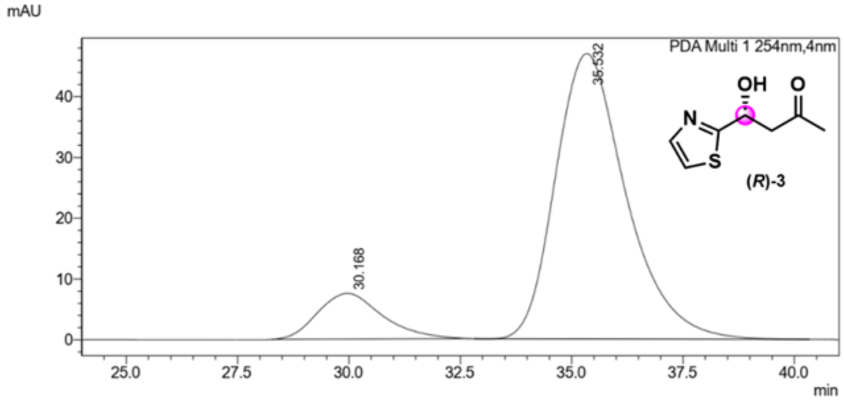


<Peak Table>
PDA Ch1 254nm

| Peak# | Ret. Time | Area | Area% |
|---|---|---|---|
| 1 | 30.168 | 763244 | 12.879 |
| 2 | 35.532 | 5163186 | 87.121 |
| Total | | 5926430 | 100.000 |

DERA$_{Ec}$-A203G

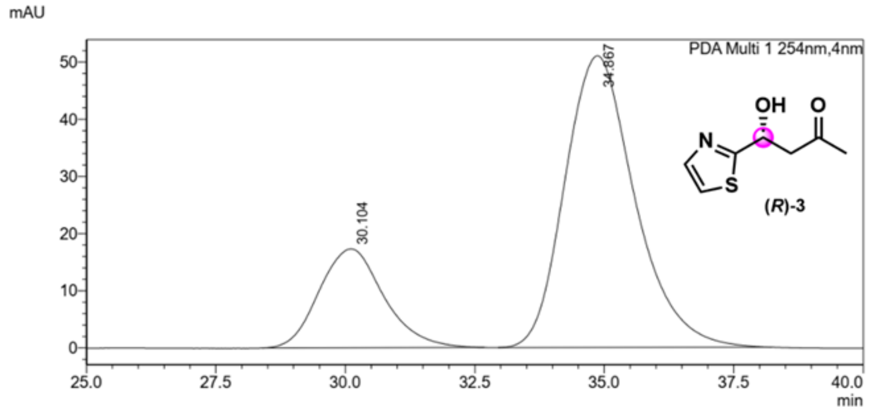


<Peak Table>
PDA Ch1 254nm

| Peak# | Ret. Time | Area | Area% |
|---|---|---|---|
| 1 | 30.104 | 1509043 | 23.913 |
| 2 | 34.867 | 4801453 | 76.087 |
| Total | | 6310497 | 100.000 |

DERA$_{Ec}$

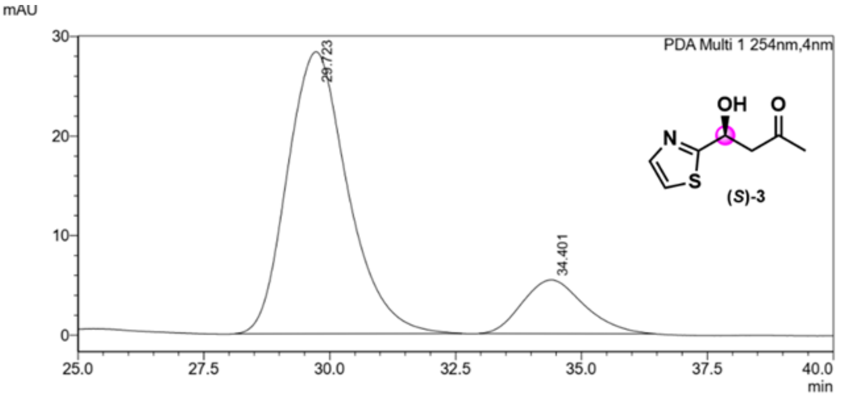


**<Peak Table>**

PDA Ch1 254nm

| Peak# | Ret. Time | Area | Area% |
|---|---|---|---|
| 1 | 29.723 | 2363343 | 83.697 |
| 2 | 34.401 | 460350 | 16.303 |
| Total | | 2823693 | 100.000 |

(14) Huang, S. K.; Picard, L.-P.; Rahmatullah, R. S. M.; Pandey, A.; Van Eps, N.; Sunahara, R. K.; Ernst, O. P.; Sljoka, A.; Prosser, R. S. Mapping the Conformational Landscape of the Stimulatory

Heterotrimeric G Protein. *Nat. Struct. Mol. Biol.* **2023**, *30* (4), 502–511. https://doi.org/10.1038/s41594-023-00957-1
(15) Bussi, G.; Donadio, D.; Parrinello, M. Canonical Sampling through Velocity Rescaling. *J. Chem. Phys.* **2007**, *126* (1), 014101. https://doi.org/10.1063/1.2408420
(16) Bernetti, M.; Bussi, G. Pressure Control Using Stochastic Cell Rescaling. *J. Chem. Phys.* **2020**, *153* (11), 114107. https://doi.org/10.1063/5.0020514
(17) Van Der Spoel, D.; Lindahl, E.; Hess, B.; Groenhof, G.; Mark, A. E.; Berendsen, H. J. C. GROMACS: Fast, Flexible, and Free. *J. Comput. Chem.* **2005**, *26* (16), 1701–1718. https://doi.org/10.1002/jcc.20291
(18) Guterres, H.; Park, S.-J.; Cao, Y.; Im, W. CHARMM-GUI *Ligand Designer* for Template-Based Virtual Ligand Design in a Binding Site. *J. Chem. Inf. Model.* **2021**, *61* (11), 5336–5342. https://doi.org/10.1021/acs.jcim.1c01156
(19) Lee, H. S.; Im, W. G-LoSA: An Efficient Computational Tool for Local Structure-centric Biological Studies and Drug Design. *Protein Sci.* **2016**, *25* (4), 865–876. https://doi.org/10.1002/pro.2890
(20) Heine, A.; Luz, J. G.; Wong, C.-H.; Wilson, I. A. Analysis of the Class I Aldolase Binding Site Architecture Based on the Crystal Structure of 2-Deoxyribose-5-Phosphate Aldolase at 0.99 Å Resolution. *J. Mol. Biol.* **2004**, *343* (4), 1019–1034. https://doi.org/10.1016/j.jmb.2004.08.066
(21) Michaud-Agrawal, N.; Denning, E. J.; Woolf, T. B.; Beckstein, O. MDAnalysis: A Toolkit for the Analysis of Molecular Dynamics Simulations. *J. Comput. Chem.* **2011**, *32* (10), 2319–2327. https://doi.org/10.1002/jcc.21787
(22) Humphrey, W.; Dalke, A.; Schulten, K. VMD: Visual Molecular Dynamics. *J. Mol. Graph.* **1996**, *14* (1), 33–38. https://doi.org/10.1016/0263-7855(96)00018-5
(23) Nosé, S. A Molecular Dynamics Method for Simulations in the Canonical Ensemble. *Mol. Phys.* **2002**, *100* (1), 191–198. https://doi.org/10.1080/00268970110089108
(24) Hoover, W. G. Canonical Dynamics: Equilibrium Phase-Space Distributions. *Phys. Rev. A* **1985**, *31* (3), 1695–1697. https://doi.org/10.1103/PhysRevA.31.1695
(25) Parrinello, M.; Rahman, A. Polymorphic Transitions in Single Crystals: A New Molecular Dynamics Method. *J. Appl. Phys.* **1981**, *52* (12), 7182–7190. https://doi.org/10.1063/1.328693
(26) Naik, K.; Jeevani, K.; Bar, K.; Saravanan, T. Enantiodivergent Synthesis of Both ( *R* )- and ( *S* )-Heteroaryl Aldols by Rationally Engineered Aldolases. *J. Org. Chem.* **2025**, acs.joc.4c02819. https://doi.org/10.1021/acs.joc.4c02819
(27) Jennewein, S.; Schürmann, M.; Wolberg, M.; Hilker, I.; Luiten, R.; Wubbolts, M.; Mink, D. Directed Evolution of an Industrial Biocatalyst: 2-Deoxy-D-Ribose 5-Phosphate Aldolase. *Biotechnol. J.* **2006**, *1* (5), 537–548. https://doi.org/10.1002/biot.200600020
(28) Rouvinen, J.; Andberg, M.; Pääkkönen, J.; Hakulinen, N.; Koivula, A. Current State of and Need for Enzyme Engineering of 2-Deoxy-D-Ribose 5-Phosphate Aldolases and Its Impact. *Appl. Microbiol. Biotechnol.* **2021**, *105* (16–17), 6215–6228. https://doi.org/10.1007/s00253-021-11462-0
